# Predisposition to Proinsulin Misfolding as a Genetic Risk to Diet-Induced Diabetes

**DOI:** 10.1101/2021.06.01.446633

**Authors:** Maroof Alam, Anoop Arunagiri, Leena Haataja, Mauricio Torres, Dennis Larkin, John Kappler, Niyun Jin, Peter Arvan

## Abstract

Throughout evolution, proinsulin has exhibited significant sequence variation in both C-peptide and insulin moieties. As the proinsulin coding sequence evolves, the gene product continues to be under selection pressure both for ultimate insulin bioactivity and for the ability of proinsulin to be folded for export through the secretory pathway of pancreatic β-cells. The substitution proinsulin-R(B22)E is known to yield a bioactive insulin, although R(B22)Q has been reported as a mutation that falls within the spectrum of Mutant *INS*-gene induced Diabetes of Youth (MIDY). Here we have studied mice expressing heterozygous (or homozygous) proinsulin-R(B22)E knocked into the *Ins2* locus. Neither females nor males bearing the heterozygous mutation develop diabetes at any age examined, but subtle evidence of increased proinsulin misfolding in the endoplasmic reticulum is demonstrable in isolated islets from the heterozygotes. Moreover, males have indications of glucose intolerance and within a few week exposure to a high-fat diet, they develop frank diabetes. Diabetes is more severe in homozygotes, and the development of disease parallels a progressive heterogeneity of β-cells with increasing fractions of proinsulin-rich/insulin-poor cells, as well as glucagon-positive cells. Evidently, sub-threshold predisposition to proinsulin misfolding can go undetected, but provides genetic susceptibility to diet-induced β-cell failure.

## Introduction

Several groups have been pursuing the molecular mechanism(s) underlying the β-cell dysfunction and compensatory cellular response(s) in the rare genetic syndrome of Mutant *INS*-gene induced Diabetes of Youth (MIDY) (1-3). The fundamental defect in MIDY is observed in humans (4), large animal models (5), small animal models (6), and has been replicated in cell culture (7), in vitro (8) and modeled in silico (9). The clinical problem originates from the fact that misfolded proinsulin within the endoplasmic reticulum (ER) can propagate its misfolding and ER retention onto wild-type (WT) bystander proinsulin molecules, thereby impairing insulin production (10; 11). And yet MIDY is a rare disease (12; 13).

Proinsulin exhibits significant sequence variation throughout evolution, including within both the C-peptide and insulin moieties (14; 15). The *Ins* gene product continues to be under selection pressure both for the ultimate bioactivity (of insulin) and for the ability of proinsulin to be folded for export through the secretory pathway of pancreatic β-cells (16). Recent evidence suggests that, with only natural variation provided by evolution, “WT” proinsulin itself is capable of forming non-native disulfide-linked proinsulin complexes not unlike those triggered by MIDY proinsulin mutations (17). However, massive formation of disulfide-linked complexes of WT proinsulin has thus far been observed only in the islets of *db/db* or other leptin receptor deficient mice [or in the islets of normal animals that have been treated with one or more toxins that drastically perturb ER homeostasis (17)]. We have also wondered whether it is possible that subtle predisposition to proinsulin misfolding can go unnoticed and yet be a genetic risk to diet-induced diabetes.

With these considerations in mind, we have interest in proinsulin residue-46, i.e., Arg at position-22 of the insulin B-chain. Recent studies indicate that insulin-R(B22)Q is a naturally-occuring variant in bats (15), has ≥ half of the normal affinity for insulin receptor binding, and can be found released into the bloodstream of patients who express the *INS* c.137G>A (R46Q) variant (18). And yet, in all three family members bearing this heterozygous mutation, diabetes was ultimately diagnosed at ages 17 – 20 years old (19), suggesting that this substitution — albeit less severe than MIDY mutants triggering neonatal diabetes (12) — still trips over the diabetogenic threshold.

It has been reported that modification of R(B22) with a bulky group of the opposite charge does not alter the specific bioactivity of insulin (20) as in the case of R(B22)D substitution (21), which is naturally occurring in mole rats and guinea pigs (15); similarly, the R(B22)E substitution has also been reported to retain insulin bioactivity (22). Here, we’ve pursued the possible impact of proinsulin-R(B22)E expression in the β-cells of mice in which this variant is knocked into the *Ins2* locus. As expected, the substitution creates a proinsulin that is secretable from pancreatic β-cells, and under normal laboratory conditions all heterozygous animals remain diabetes-free. Nevertheless, heterozygous males consistently developed diabetes upon exposure to a high-fat diet, highlighting the impact of a sub-threshold genetic predisposition to proinsulin misfolding on the development of diet-induced diabetes.

## Materials and Methods

### Proinsulin mutagenesis

Plasmids encoding untagged human proinsulin-R(B22)E, untagged human proinsulin-WT, or Myc-tagged proinsulins were generated as described (23; 24). All proinsulin-expressing constructs were confirmed by direct DNA sequencing.

### Cell transfection, metabolic labeling, immunoprecipitation (IP), co-IP, Western blotting

Min6 mouse β-cells (25) (obtained Dr. D. Stoffers, U. Pennsylvania) were cultured in DMEM supplemented with 10% FBS, penicillin/streptomycin and 0.05 mM β-mercaptoethanol. Cells at 70-80% confluency were transfected using Lipofectamine 2000 (Thermo-Fisher) with fresh media changed at 6-h post-transfection. Media were removed at 48 h; the cells were washed with ice-cold PBS and lysed in RIPA buffer (10 mM Tris pH 7.4, 150 mM NaCl, 0.1% SDS, 1% NP40, 2 mM EDTA) plus protease inhibitor/phosphatase inhibitor cocktail (Sigma-Aldrich). Total protein was measured by BCA. Proteins in sample buffer were boiled and resolved by 4–12% Bis-Tris NuPAGE (Invitrogen) and electrotransferred to nitrocellulose (Bio-Rad). Membranes were incubated with primary antibodies (4°C overinight) and then secondary peroxidase-conjugated Goat Anti-Rabbit IgG (Jackson ImmunoResearch 111-035-144) or peroxidase-conjugated Goat Anti-Mouse IgG (Jackson Immuno Research 115-035-174, followed by ECL (SuperSignal West Pico PLUS, Thermo-Fisher) with digital image capture.

### Construction of Knock-in mouse

Initially, the Mouse Genetics Core (National Jewish Health) prepared eggs from superovulated NOD females, fertilized with male NOD sperm *in vitro*. After overnight culture, fertilized embryos were injected with a mixture of: 1) Cas9 protein, 2) a CRISPR guide RNA (designed using CRISPOR software, http://crispor.tefor.net/) matching a region just upstream of the proinsulin R(B22) codon, and 3) an *Ins2*-specific 150 bp repair oligonucleotide covering this region replacing the R codon (CGT) with E (GAG) plus a silent mutation creating an *Alu I* restriction site. Injected embryos were introduced into pseudo-pregnant mice; DNA from the pups were analyzed by PCR, restriction digest, and sequencing to distinguish specific *Ins2-* (or offsite *Ins1-*) locus repair with the homologous sequence versus non-homologous end-joining events. To isolate the proper genetic event, breeding resulted in NOD mice bearing a single heterozygous *Ins2* R(B22)E mutation (WT at three other *Ins* alleles). Breeding was then initiated to move the mutation into the C57BL6/j background. Animals were phenotyped in each of the first 5 generations of C57BL6/j backcrosses. All data from this manuscript come after 5 generations of backcrossing (and further backcrosses are ongoing), but as the same phenotype was observed each backcross generation, the data are reported here. High-fat diet (from 5.5 – 11.5 weeks of age) was irradiated rodent chow (60 kcal% fat; #D12492, Research Diets, New Brunswick NJ).

### Circulating insulin and proinsulin; *in vivo* glucose tolerance; glucose-stimulated insulin secretion (GSIS)

Mouse insulin: ALPCO ELISA, 80-INSMS-E10; and proinsulin: ALPCO ELISA, 80-PINMS-E01. For *in vivo* glucose tolerance, mice were fasted for 6 h; glucose (1g/kg body weight) was administered intraperitoneally; tail vein glucose was monitored (One-Touch Ultra meter and test strips). For *in vivo* GSIS, serum was collected under basal conditions and under glucose-stimulated conditions as at t=15 min.

### Islet isolation and GSIS

Islets were isolated by collagenase digestion via the common bile duct, followed by pancreatic digestion *ex vivo*. The digest was washed and spun on a Histopaque (1077-Sigma-Aldrich) gradient (900 *g* x 20 min without brake). Islets were collected washed, handpicked, and incubated overnight in RPMI-1640 medium plus 10% FBS, at 37°C. Recovered islets were pre-incubated at 2.8 mM glucose for 1 h at 37°C in modified Krebs-Ringer bicarbonate buffer plus 20 mM HEPES and 0.05% BSA (KRBH-BSA). 15-17 islets were transferred to microfuge tubes containing 500 uL KRBH-BSA solution and incubated at 37°C for 30 min a 2.8 mM glucose (basal) followed by 30 min at 16.7 mM glucose (stimulated), with media measured for insulin content.

### Metabolic labeling of mouse pancreatic islets

25 islets isolated from WT and proinsulin-R(B22)E heterozygous and homozygous littermates were washed in prewarmed Met/Cys-deficient RPMI medium and then pulse-labeled with ^35^S-amino acids (Tran^35^S label) for 30 min at 37°C. Labeled islets were either lysed immediately, or chased in complete growth media for 2h. Islets were sonicated in RIPA buffer (25 mM Tris, pH 7.5, 100 nM NaCl, 1% Triton X-100, 0.2% deoxycholic acid, 0.1% SDS, 10 mM EDTA) containing 2 mM NEM and a protease inhibitor cocktail. Lysates were normalized to trichloroacetic acid-precipitable counts, and immunoprecipitated with guinea pig polyclonal anti-insulin and protein A-agarose overnight at 4°C. Immunoprecipitates were washed and analyzed by nonreducing / reducing Tris– tricine–urea–SDS-PAGE, followed by phosphorimaging, and bands quantified with ImageJ software.

### Immunofluorescence

Paraffin sections of formaldehyde-fixed pancreas were de-paraffinized with Citrisolv (Fisher Scientific), re-hydrated in a decreasing graded series of ethanol, followed by heating for antigen retrieval. Slides were washed with PBS and incubated in blocking buffer (TBS plus 0.2% Triton X-100 and 3% BSA) for 2 h, and incubated in primary antibody (in TBS plus 3% BSA, 0.2% Tween-20) overnight at 4 °C. After washes, secondary antibody was incubated for 1 h at RT. Slides were washed thrice with TBS/0.1% Tween-20, mounted with Prolong Gold + DAPI, and epifluorescence imaged on a NikonA1 confocal microscope with a 60x oil objective.

### Electron Microscopy

Isolated islets were fixed with 2.5% (v/v) glutaraldehyde in 0.1 M sodium cacodylate buffer pH 7.2 (CB), embedded in 2.5% low melting agarose, trimmed to ∼1mm cubes, washed 3 times in CB, and post-fixed in 2% osmium tetroxide plus 1.5% potassium ferrocyanide in 0.1M CB for 1 h on ice. After 3 further washes in CB, cubes were washed thrice in 0.1M sodium acetate buffer pH 5.2 and then stained in this buffer containing 2% uranyl acetate for 1 h. With agitation the cubes were washed extensively and then dehydrated in a graded series of ethanol up to 100% and then acetone for 15 min, before infiltration with graded concentrations of Spurr’s resin in acetone over 3 d. Islets were finally embedded in fresh Spurr’s resin and polymerized for 45 h at 70-75°C. Ultra-thin (70 nm) sections were captured on carbon-coated 200-mesh copper grids, and imaged on a JEOL JEM-1400-plus transmission electron microscope (JEOL USA Inc., Peabody, MA) at 80kV, with images captured on an AMT XR401 camera (Advanced Microscopy Techniques, Woburn, MA).

### Statistical analyses

Statistical analyses were carried out by two-tailed student’s t-test or one-way ANOVA followed by Dunnett multiple comparisons using GraphPad Prism 8. A *p*-value of < 0.05 was taken as statistically significant.

### Data and resource availability

Data generated and analyzed during the current study are contained within the Figures; additional data are available from the corresponding author upon request.

## Results

### Expression of recombinant proinsulin-R(B22)E

Proinsulin-R(B22)E (Fig. 1A) should lead to bioactive insulin (20-22), yet may not necessarily support efficient proinsulin protein folding (16; 26-29). We first examined the behavior of recombinant human proinsulin-R(B22)E. Min6 pancreatic β-cells express their own endogenous mouse proinsulin, but we probed transfected cells, and media, with an antibody that recognizes only human proinsulin protein. Thus, from Min6 cells transfected with empty vector, no proinsulin band was detected (Fig. 1B). Both human WT and R(B22)E proinsulins were secreted from Min6 cells (Fig. 1B), and under these conditions, proinsulin-R(B22)E did not block WT proinsulin secretion (Fig. 1C). Nevertheless, from Western blotting of reducing SDS-PAGE, it was clear that the secretion efficiency (extracellular-to-intracellular ratio) for WT proinsulin was greater than that for proinsulin-R(B22)E, whereas *Akita* mutant proinsulin was not secreted at all (Fig. 1B, C). Thus, in Min6 β-cells, proinsulin-R(B22)E passes ER quality control, and a dominant-negative MIDY phenotype is inapparent.

**Fig. 1.**
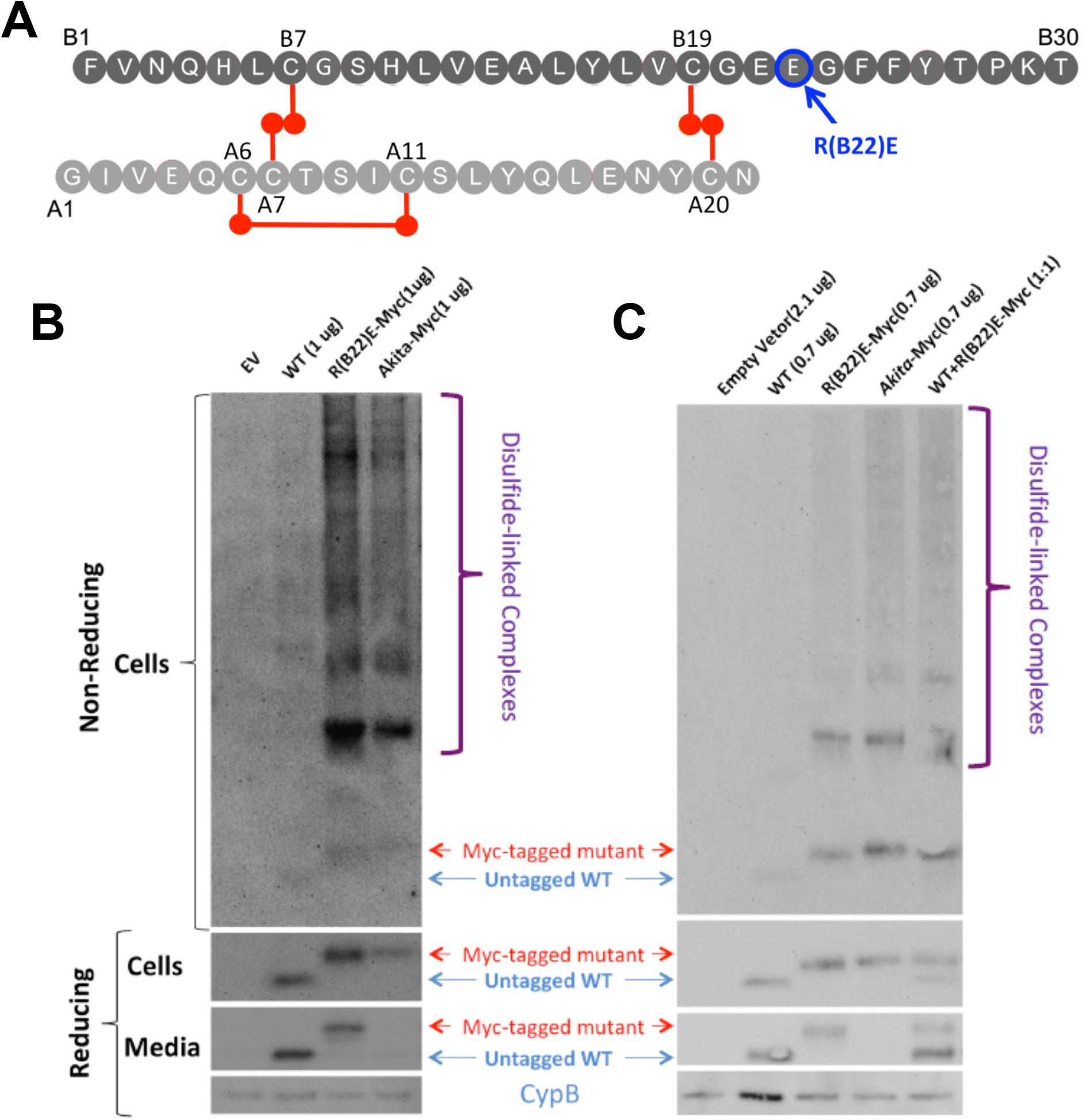
Insulin B-chain substitution R(B22)E. **A)** Schematic representation of insulin chains showing the respective disulfide bonds and the R(B22)E substitution. **B)** Transfection of Min6 cells with empty vector (EV), untagged wild-type (WT) human proinsulin, hPro-R(B22)E-CpepMyc, or hPro-C(A7)Y-CpepMyc (‘Akita-Myc’). The media bathing transfected cells was collected overnight, and both cell lysates and media were resolved by SDS-PAGE under reducing conditions and immunoblotting with anti-human-proinsulin or cyclophilin B (CypB, loading control) in the panels below. Cell lysates were similarly analyzed under nonreducing conditions to reveal disulfide-linked proinsulin complexes, in the upper panel. **C)** Transfection of Min6 cells as in panel B (first four lanes) or co-transfection with both untagged WT human proinsulin and hPro-R(B22)E-CpepMyc (last lane). Proinsulin-R(B22)E is secreted from Min6 cells, and under these conditions, proinsulin-R(B22)E does not block the secretion of co-expressed untagged WT proinsulin.

### Endogenous islet expression of *Ins2*-proinsulin-R(B22)E

A CRISPR/Cas9-mediated knock-in of *Ins2*-proinsulin-R(B22)E was back-bred to C57BL6/j mice. From 5.5 to 11.5 weeks of age, WT and heterozygous *Ins2*-proinsulin-R(B22)E mice gained weight on a normal chow diet, and as expected, gained more weight on a high-fat diet (**HFD**), without statistical differences between the genotypes (Fig. 2A). Glucose tolerance in heterozygous *Ins2*-proinsulin-R(B22)E mice at 5.5 weeks of age was normal (Fig. 2B) with normal area under the curve (Fig. 2C). However, within 2 weeks of HFD, male heterozygotes had an average random blood glucose ≥ 350 mg/dL, suggesting onset of diabetes (Fig. 2D). At 11.5 wks of age — either in terms of fasting hyperglycemia, 2-hr glucose tolerance, or area under the curve — male heterozygotes on HFD for 6 weeks had clearly diagnosed diabetes, whereas those on a normal chow diet did not meet any diabetes criteria (Fig. 2E, F). Despite that random glucose in HFD-fed male heterozygotes was higher, serum insulin was not elevated (Fig. 2G); actually, the random serum insulin-to-glucose ratio was significantly lower in HFD-fed male heterozygotes (Fig. 2H). Indeed, after 6 weeks of HFD, isolated islets from male *Ins2*-proinsulin-R(B22)E heterozygotes exhibited abnormally low insulin secretion (unstimulated and glucose-stimulated) despite a normal fold-change (Fig. 2I). Islets from male heterozygotes also developed diminished insulin content and in HFD-fed animals insulin content fell further (Fig. 2J). Additionally, proinsulin in these mice formed enhanced non-native disulfide-linked complexes (Supplement S1). In contrast, female *Ins2*-proinsulin-R(B22)E heterozygotes gained weight normally and did not even develop random hyperglycemia on HFD (Fig. 2K, L).

**Fig. 2.**
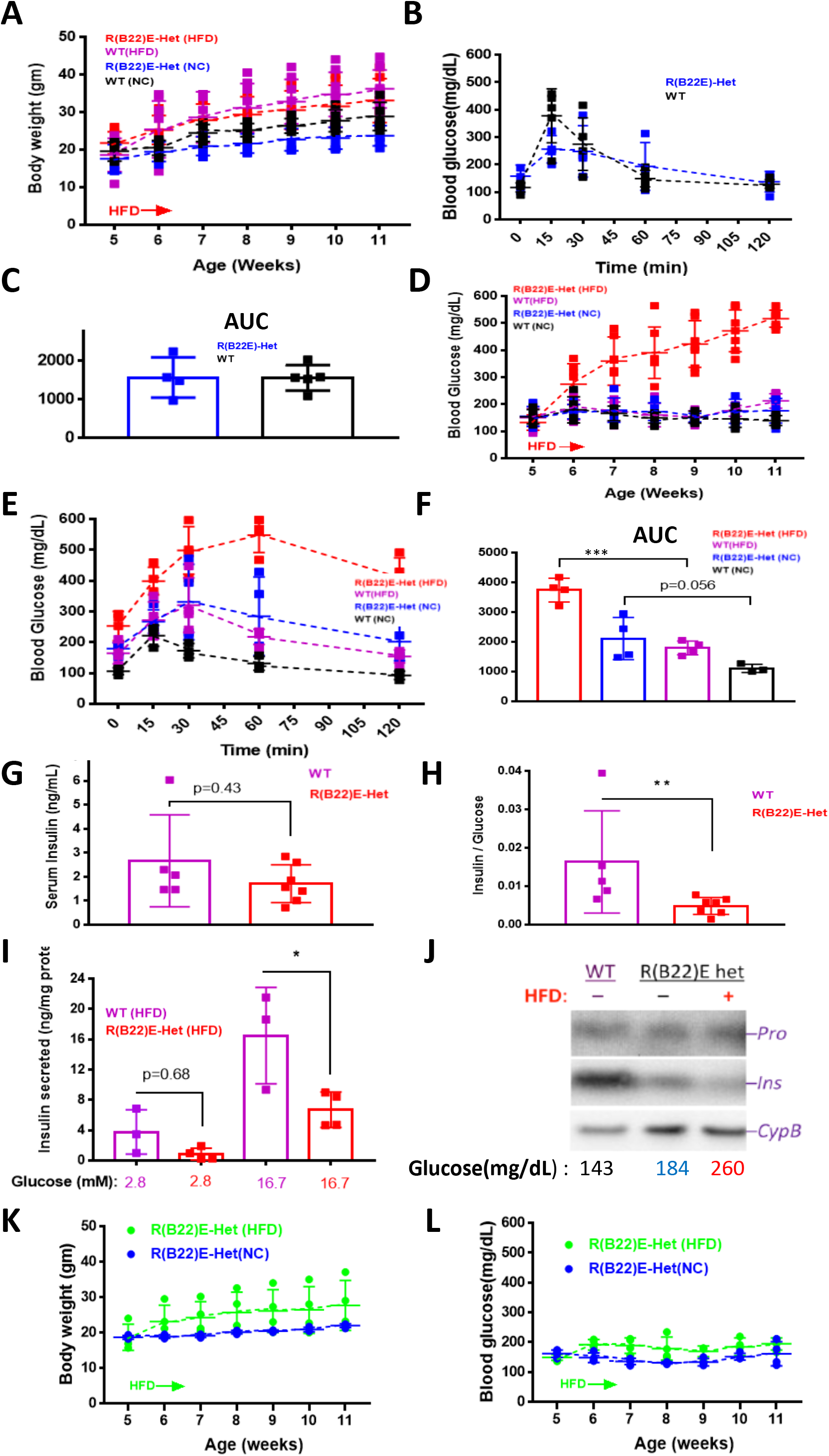
Heterozygous *Ins2*-proinsulin-R(B22)E mice fed normal chow (NC) or high fat diet (HFD). **A)** Weekly body weight measurements in male mice (n = 4-8 per group). **B)** Intraperitoneal glucose tolerance test (IPGTT) in 5.5-week-old males on normal chow (*n* = 4-5 per group). **C)** Area under the curve (AUC) from the data in panel B. **D)** Weekly random blood glucose in males on normal chow diet or HFD [n = 4-8 per group; *Ins2*-proinsulin-R(B22)E heterozygotes on normal diet in blue, and on HFD in red]. **E)** IPGTT in 11 week-old males on normal chow diet or HFD for 6 weeks [n = 3-5 per group; *Ins2*-proinsulin-R(B22)E heterozygotes on normal diet in blue, and on HFD in red]. **F)** AUC from the data in panel E. **G)** Random serum insulin level in 11 week-old WT (purple bar) or *Ins2*-proinsulin-R(B22)E heterozygous males (red bar) on HFD for 6 weeks (n = 5-7 per group). **H)** The data from panel G used to calculate serum insulin-to-glucose ratio. **I)** Insulin secretion from isolated islets of WT (purple bars) or heterozygous *Ins2*-proinsulin-R(B22)E heterozygous males on HFD (red bars) at 2.8 mM (unstimulated) and 16.7 mM (stimulated) glucose (n = 3-4 per group). **J)** Isolated islets from male WT and heterozygous *Ins2*-proinsulin-R(B22)E mice fed normal chow or HFD were lysed and analyzed by reducing SDS-PAGE and immunoblotting with mAb anti-proinsulin (upper panel), guinea pig anti-insulin (middle panel), and anti-cyclophilin B (CypB, loading control). **K)** Weekly body weight measurements in female heterozygous *Ins2*-proinsulin-R(B22)E mice fed normal chow (blue) or HFD (green) (n = 4 per group). **L)** Weekly random blood glucose measurements in female heterozygous *Ins2*-proinsulin-R(B22)E mice fed normal chow (blue) or HFD (green) (n = 4 per group). All graphs include mean ± SD. \**p* < 0.05; \*\**p* < 0.01; \*\*\**p* < 0.001; 1-way ANOVA.

When *Ins2*-proinsulin-R(B22)E was bred to homozygosity (still leaving both WT *Ins1* alleles), even on a normal chow diet, at 5 weeks of age both males and females exhibited random blood glucose that averaged in the diabetic range (Fig. 3A) and fasting blood glucose values were similarly elevated. [Random blood glucose in homozygotes rose even higher by 8 weeks of age, whereas heterozygotes remained normoglycemic (Fig. 3B).] Despite the higher blood glucose, neither homozygous males nor females raised endogenous serum insulin; with both sexes combined it was apparent that homozygotes had decreased insulinemia despite ongoing hyperglycemic stimulation (Fig. 3C), with a profound lowering of serum insulin-to-glucose ratio (Fig. 3D). These three genotypes were clearly separable when comparing acute GSIS *in vivo* (Fig. 3E). Moreover, isolated islets from these subgroups tested for GSIS *in vitro* were distinct (Fig. 3F), despite that the *fold-change* of GSIS was not statistically different. The data indicate a decline of releasable insulin beginning with *Ins2*-proinsulin-R(B22)E heterozygotes, and progressively worsening in homozygotes.

**Fig. 3.**
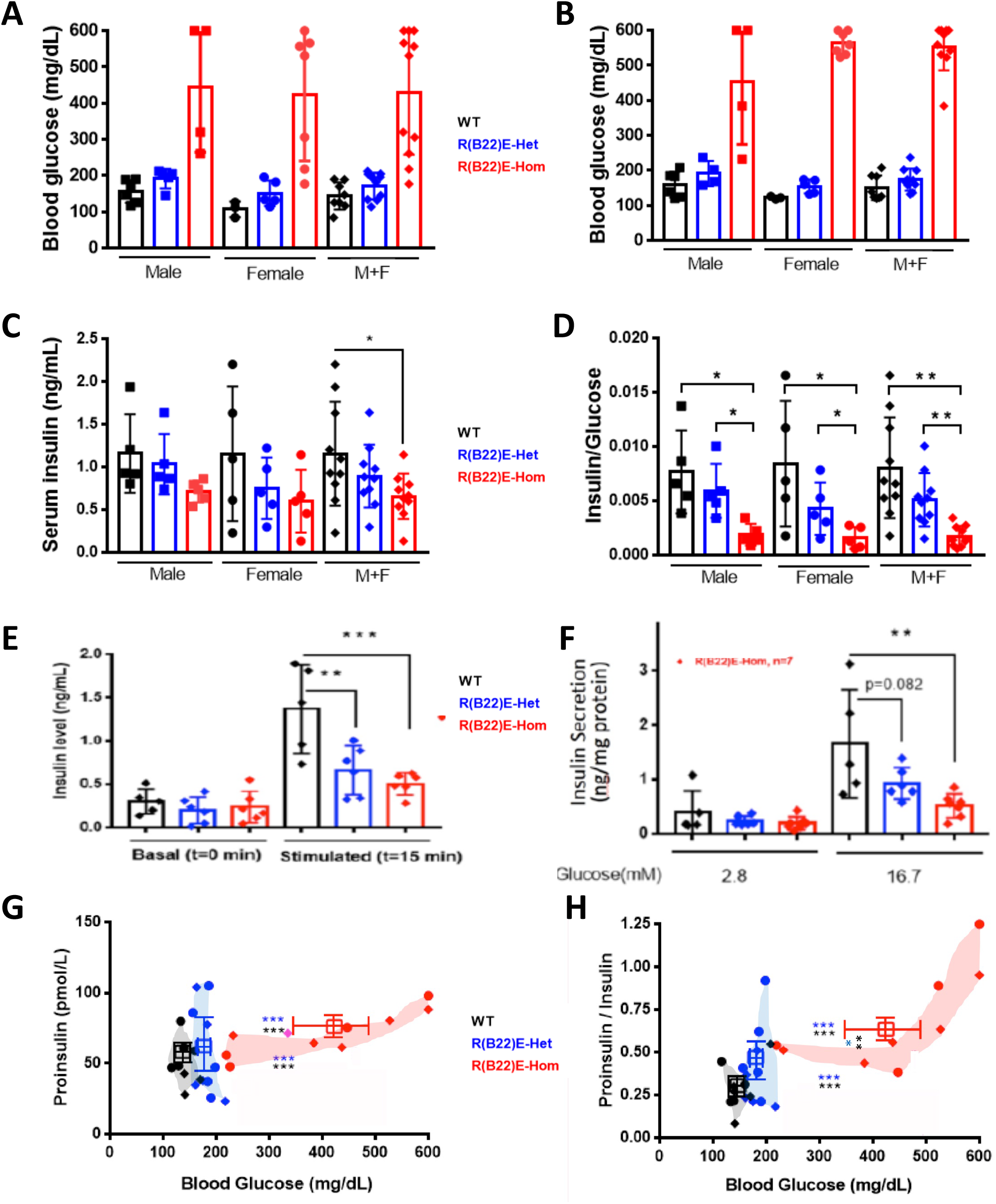
Homozygous *Ins2*-proinsulin-R(B22)E mice develop spontaneous diabetes. **A)** Random blood glucose at 5 weeks age in WT (black bars), heterozygous (blue bars), and homozygous (red bars) *Ins2*-proinsulin-R(B22)E mice fed normal chow (n = 3-7 per group). Males and females shown separately as well as combined together. **B)** The same mice from panel A re-measured 8 weeks postpartum. **C)** Serum insulin level measured at 5.5 weeks of age in WT (black bar), heterozygous (blue bars), or homozygous (red bars) males fed normal chow (n = 5 per group). Males and females shown separately as well as combined together. **D)** Insulin-to-glucose ratio calculated from the samples in panels B-C, respectively. **E)** GSIS i*n vivo* at 0 and 15 min post-stimulation from WT (black bars), heterozygous (blue bars), or homozygous (red bars) *Ins2*-proinsulin-R(B22)E mice fed normal chow (6 week-old males; n = 5-6 per group). **F)** GSIS from islets isolated from WT (black bars), heterozygous (blue bars), or homozygous (red bars) *Ins2*-proinsulin-R(B22)E mice fed normal chow, at 2.8 mM (unstimulated) and 16.7 mM (stimulated) glucose concentration (6 week-old males; n = 5-7 per group). **G)** Serum proinsulin level from the same animals measured in panel C (males and females combined together). **H)** Proinsulin-to-insulin ratio from the animals in panel G (males and females combined together). All graphs include mean ± SD. \**p* < 0.05; \*\**p* < 0.01; \*\*\**p* < 0.001; 1-way ANOVA.

Porte and Kahn, as well as others — reported that patients with type 2 diabetes demonstrate disproportionately elevated levels of circulating proinsulin that tends to be worse with the degree of hyperglycemia and correlates with a diminished maximal insulin secretion capacity (30; 31); yet recent reports suggest that serum proinsulin-to-insulin ratio measurements may be of limited value across populations (32; 33) although they may still be of value within selected subgroups (33). As model organisms have reduced genetic heterogeneity, we observed in normal chow-fed animals at 5.5 weeks of age, male+female mice solely expressing WT proinsulin (black symbols), *Ins2*-proinsulin-R(B22)E heterozygotes (blue symbols) and homozygotes (red symbols) tended to fall in three distinct groups when plotting circulating proinsulin levels versus simultaneous random blood glucose (Fig. 3G). *Ins2*-proinsulin-R(B22)E heterozygotes showed the greatest variability of circulating proinsulin amongst individuals, which was not well-correlated with random glucose, and overall these values were nearly normal (Fig. 3G). However, homozygotes formed a discrete group suggesting a positively-sloped relationship of circulating proinsulin with simultaneous random blood glucose; given the developing insulin deficiency (Fig. 3C), these differences were amplified when considering circulating proinsulin-to-insulin ratio in homozygotes (Fig. 3H), although there remained only small differences between WT and heterozygous animals. [A separate set of heterozygous HFD-fed males at 11 weeks of age (distinct from the animals shown in Figs. 3G, H) progressed into diabetes, yet the mean value for circulating proinsulin or proinsulin-to-insulin ratio was not increased (Supplemental Fig. S2)]. Altogether, the data support that circulating proinsulin-to-insulin ratio relative to prevailing glucose does serve to distinguish animal subgroups.

We next examined islet insulin and proinsulin content and localization in fixed tissues. In WT mice, β-cells exhibit robust insulin immunofluorescence, and proinsulin is concentrated in a juxtanuclear subregion (Fig. 4A). In *Ins2*-proinsulin-R(B22)E heterozygous males, again most islet cells are β-cells exhibiting obvious insulin immunofluorescence, but there were a few cells with diminished insulin signal and more expansive proinsulin immunofluorescence [suggesting depletion of insulin stores yet maintaining biosynthetic activity (Fig. 4B)]. Young *Ins2*-proinsulin-R(B22)E homozygotes that are not yet diabetic did not initially appear very different from heterozygotes: animals with random blood glucose < 200 mg/dL exhibited a subset of bright insulin-positive cells plus other islet cells with diminished insulin accompanied by increased proinsulin immunofluorescence (Fig. 4C). Even after homozygotes developed frank diabetes, there were still a few islet cells brightly immunofluorescent for insulin, but an enlarged fraction of cells showed diminished insulin signal while exhibiting robust proinsulin immunofluorescence — seen in both larger and smaller islets (Figs. 4D, E).

**Fig. 4.**
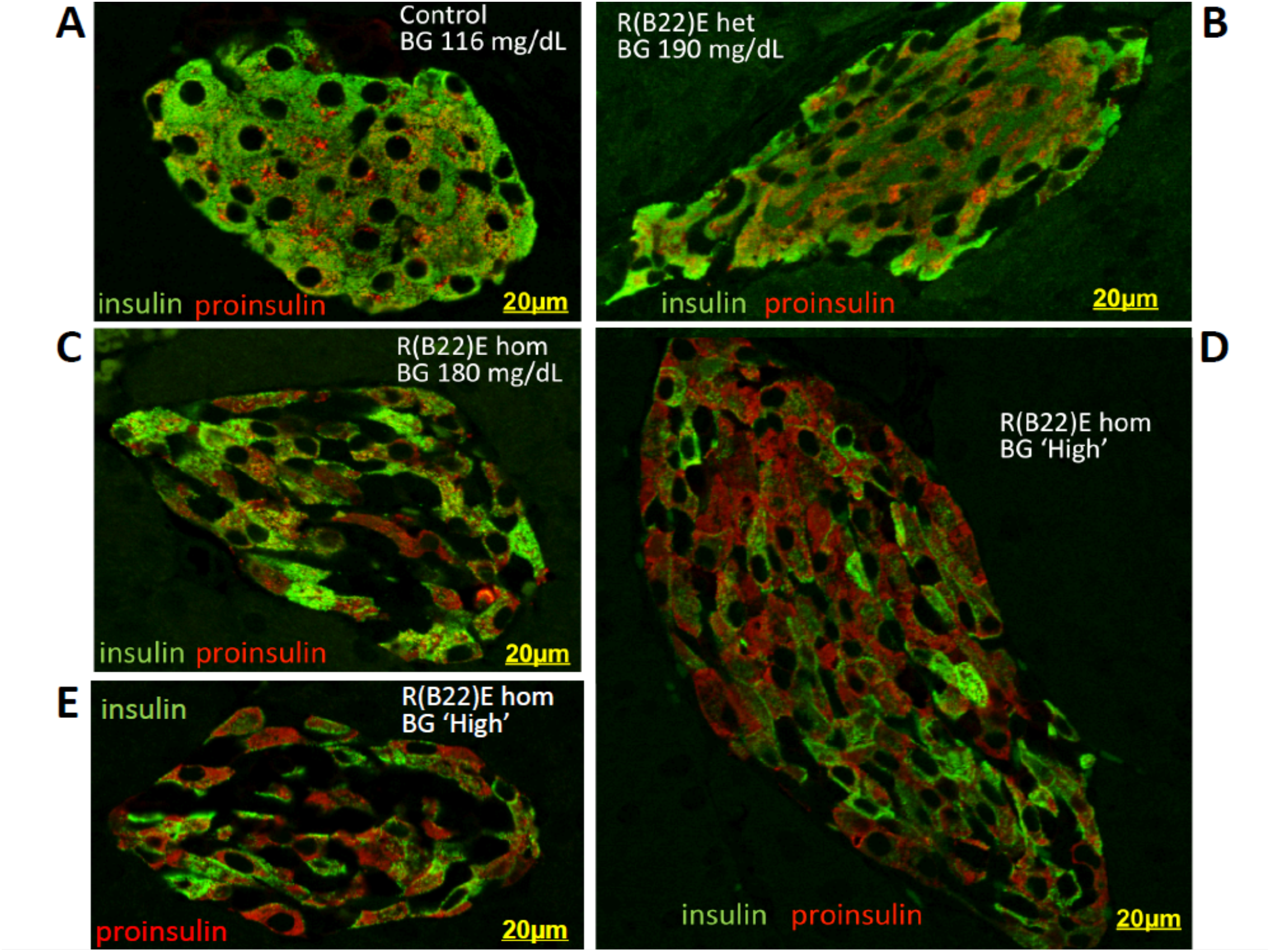
Proinsulin and insulin double-immunofluorescence. BG = random blood glucose at the time of euthanasia. **A)** WT control. **B)** *Ins2*-proinsulin-R(B22)E heterozygote (6 week-old female). **C)** Normoglycemic 4 week-old female *Ins2*-proinsulin-R(B22)E homozygote. **D and E)** *Ins2*-proinsulin-R(B22)E 7 week-old female homozygote.

By electron microscopy, β-cells from WT and normoglycemic heterozygous *Ins2*-proinsulin-R(B22)E mice had a similar, well-granulated appearance (Supplement Fig. S3A, B). Even in HFD-fed animals, the secretory pathway of WT β-cells exhibited the normal cisternal ER, ER-Golgi Vesiculo-Tubular Clusters (VTCs, pre-Golgi intermediates), well-developed Golgi stacks, as well as immature (ISGs) and mature insulin granules (Supplement S3C) — these features were also present in *Ins2*-proinsulin-R(B22)E heterozygotes on a normal chow diet (Supplement Fig. S3A, B). However on a HFD, a β-cell subpopulation in R(B22)E heterozygotes developed expanded ER and under-filled, low electron-density granule contents (“Cell B”, Supplement Fig S3D). In homozygotes, it was easy to identify a subset of β-cells bearing some insulin granules while other β-cells exhibited few granules but increased ER (Fig. 5, Supplement Fig. S3E), as well as cells bearing under-filled low-electron-density “micro-granules” and related secretory pathway organelles (Supplement Fig. S3F, G).

**Fig. 5.**
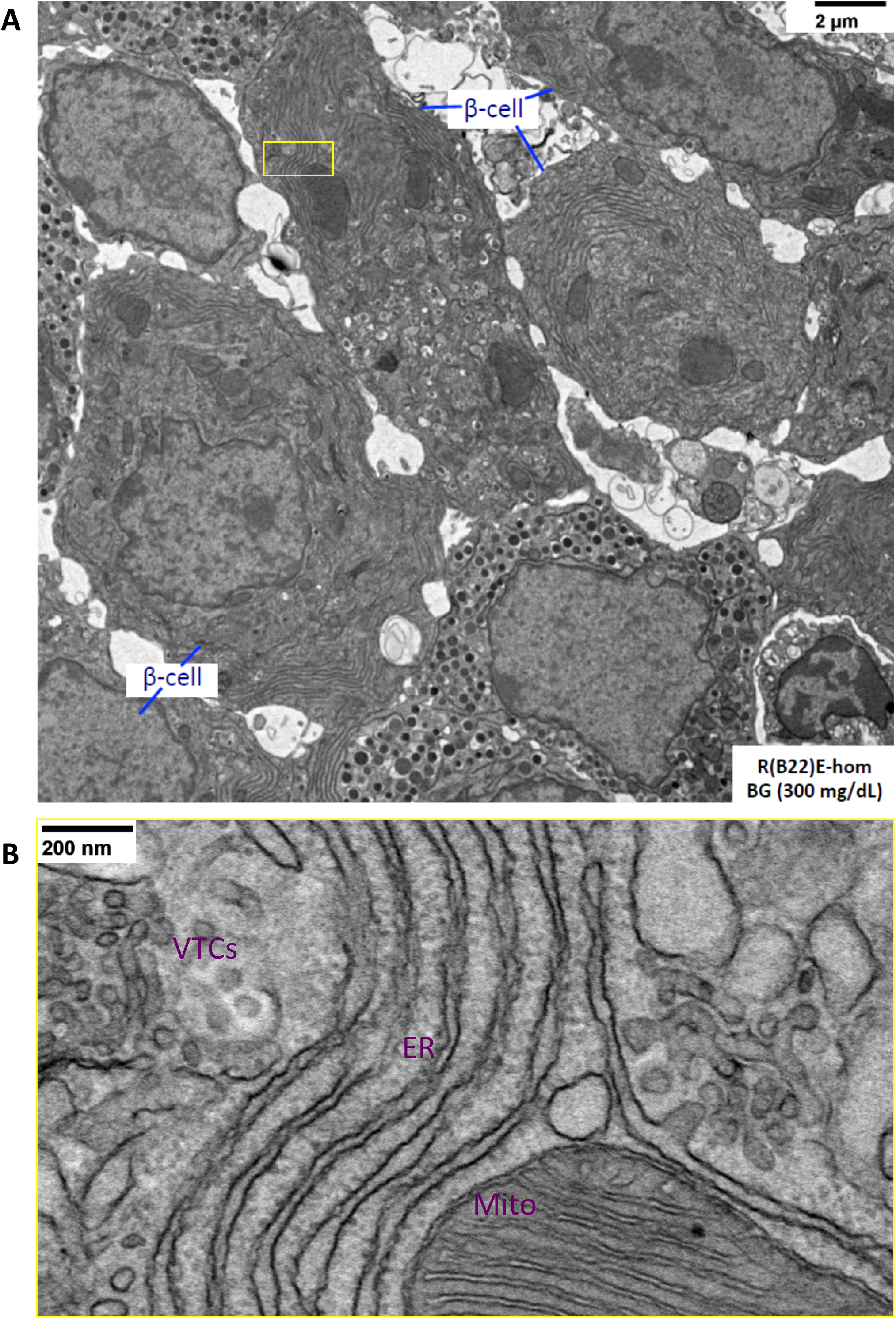
Transmission electron microscopy of islet cells from a 4 week-old male *Ins2*-proinsulin-R(B22)E homozygote, highlighting heterogeneity in the abundance of insulin secretory granules. **A)** Lower power view (scale bar in the upper right corner); β-cells are identified on the figure. The cell at bottom and two cells in the upper left corner exhibit glucagon secretory granules. BG = random blood glucose at the time of euthanasia. **B)** The yellow boxed area from panel A is shown at higher power (scale bar in the upper left corner). VTCs are a pre-Golgi compartment. Mito = mitochondrion. A more detailed electron microscopy survey is presented in the Supplement Fig. S3.

Insulin/proinsulin double-immunofluorescence images of heterozygous and homozygous *Ins2*-proinsulin-R(B22)E animals also revealed islet cells unlabeled for either marker (Fig. 4B-E). Three-color immunofluorescence in *Ins2*-proinsulin-R(B22)E-positive islets labeled glucagon-positive cells — initially at the perimeter of normoglycemic, WT islets (Fig. 6A) — which appeared increasingly within the islet interior especially in homozygous mice — and corresponded to most of the remaining cells (Fig. 6B-E). Together, the data in Figures 4-6 indicate that, progressing from WT-to-heterozygous-to-homozygous animals, a decrease of insulin-positive β-cells with increasing proinsulin-enriched cells and glucagon-enriched cells. The loss of insulin in homozygotes was also observed by immunoblotting of islet lysates analyzed by reducing SDS-PAGE. Insulin deficiency was less noticeable when the homozygotes were still at the euglycemic stage but was exacerbated in parallel with the progression of hyperglycemia (Fig. 7A+B).

**Fig. 6.**
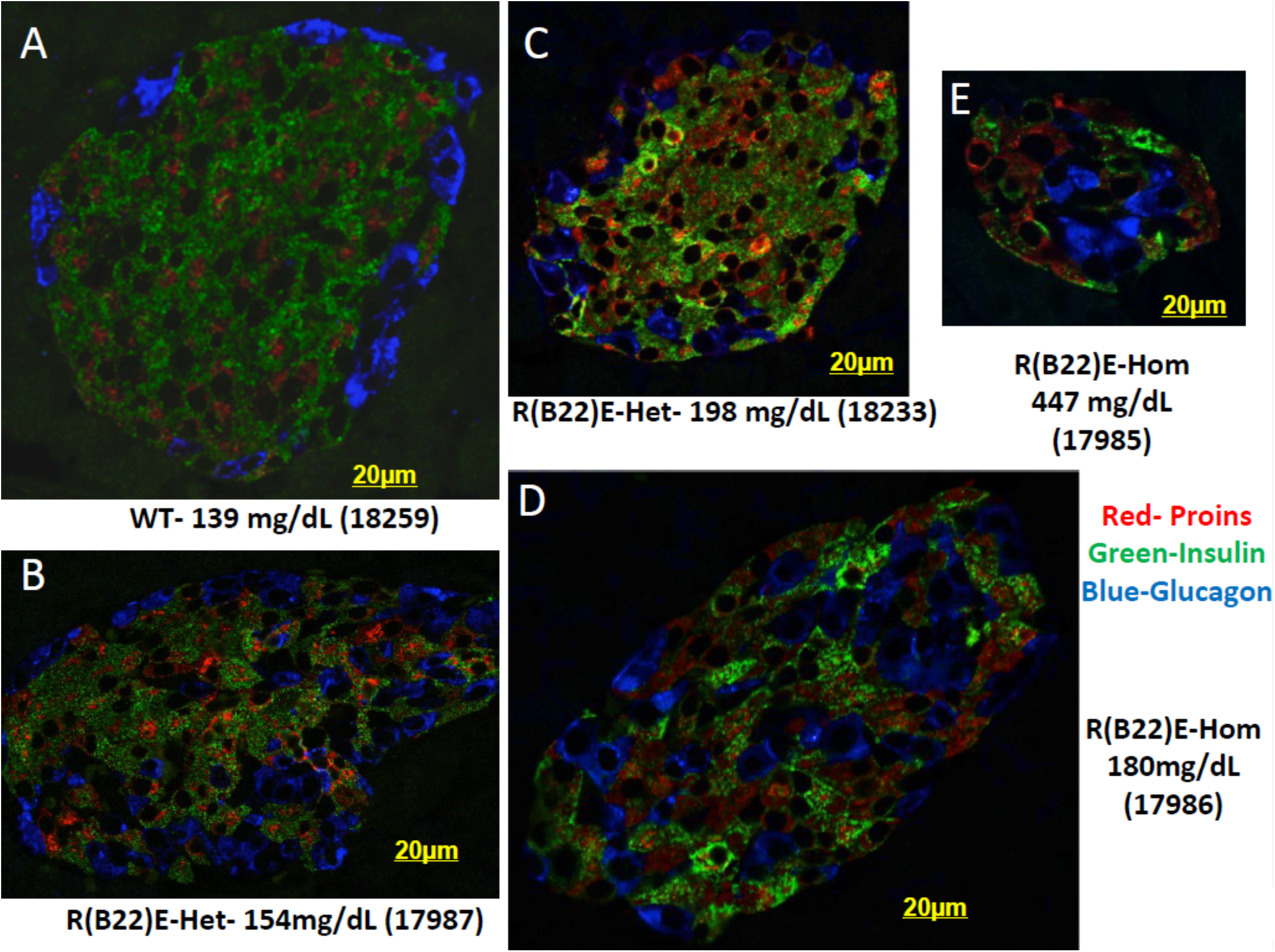
Growing subpopulations of proinsulin-enriched cells (red) and glucagon-enriched cells (blue), and a decline of insulin-enriched cells (green), in *Ins2*-proinsulin-R(B22)E heterozygotes and homozygotes, as identified by triple immunofluorescence. Random blood glucose levels are indicated on the figure. **A)** Wild-type (WT, 6 week-old female). **B and C)** *Ins2*-proinsulin-R(B22)E heterozygous females (age 6.5 weeks). **D and E)** *Ins2*-proinsulin-R(B22)E female homozygotes (ages 4 and 6 weeks, respectively).

**Fig. 7.**
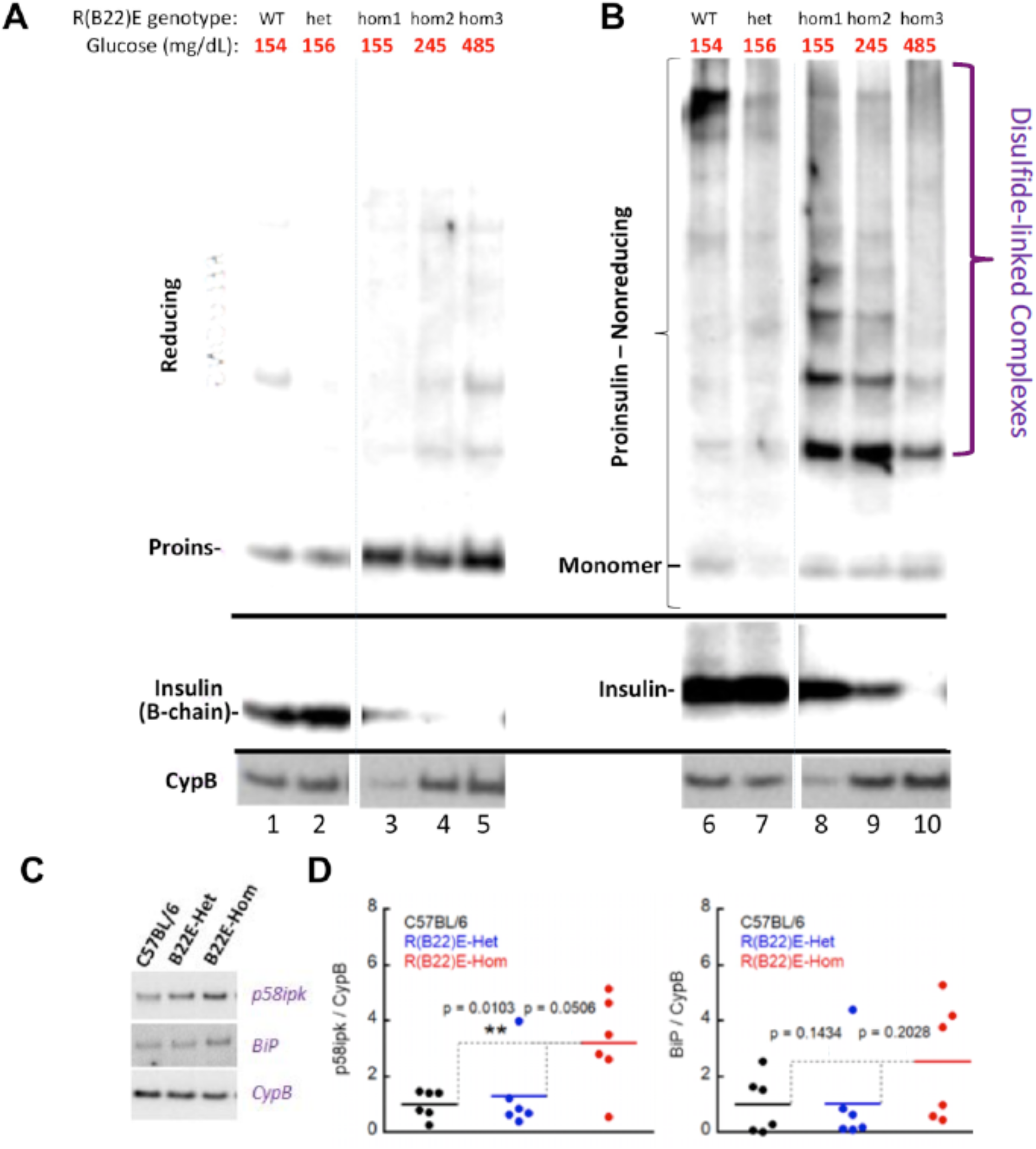
Progressive loss of insulin, accompanied by proinsulin misfolding, in *Ins2*-proinsulin-R(B22)E homozygotes. **A)** Reducing SDS-PAGE and immunoblotting with anti-proinsulin (upper panel), anti-insulin (middle panel), and anti CypB (loading control, bottom panel). Islets were isolated from males (ages 5-7 weeks) with the genotypes shown above; random blood glucose at the time of euthanasia is also indicated. **B)** Nonreducing SDS-PAGE and immunoblotting of the identical samples as that shown in panel A, highlighting aberrant proinsulin disulfide-linked complexes. **C)** Immunoblotting of p58ipk and BiP from male WT, *Ins2*-proinsulin-R(B22)E heterozygotes and homozygotes (age 5-6 weeks). **D)** Quantitation of immunoblotting like that shown in panel C (n = 6 per group); statistical analysis shown directly on the figure panels.

Remarkably, islets of *Ins2*-proinsulin-R(B22)E homozygotes did not exhibit proinsulin deficiency even as hyperglycemia progressed into the 400 – 500 mg/dL range (Fig. 7A). However, in homozygous *Ins2*-proinsulin-R(B22)E mice, a greater fraction of proinsulin was contained in aberrant disulfide-linked complexes (Fig. 7B), which have been reported both in stressed human islets and murine diabetes models (17). Indeed, even before development of frank diabetes, we noted a tendency towards increased islet BiP protein [as well as the BiP–co-chaperone, p58ipk (Fig. 7C, D)], although these effects were only ∼2-fold. In the proinsulin-enriched cells of heterozygotes and homozygotes, proinsulin extensively co-localized with ER resident proteins (reactive with anti-KDEL, Supplement Fig. S4), consistent with cellular heterogeneity that includes an increased fraction of proinsulin-enriched β-cells maintaining an expanded ER compartment.

We conducted three independent pulse-chase radiolabeling experiments to look at the efficiency of and insulin biosynthesis. First, the amount of newly-synthesized proinsulin (made in a 15 min pulse-labeling with ^35^S-amino acids) was analyzed by immunoprecipitation (IP) with anti-insulin, reducing SDS-PAGE, and autoradiography (*right half of* Fig. 8A). At 2 h of chase, cells and media were combined together before IP, and the yield of newly-made insulin (*left half of* Fig. 8A) derived from labeled proinsulin in WT control islets was nearly 100%, whereas insulin generation in euglycemic homozygous *Ins2*-proinsulin-R(B22)E islets was less efficient (*quantitation in* Supplement Fig. S5). In the next experiment, labeled heterozygous *Ins2*-proinsulin-R(B22)E islets were compared against those from a homozygote with random blood glucose of 400 mg/dL (Fig. 8B). From the euglycemic heterozygote, recovery of labeled insulin at 2h chase from the original newly-synthesized proinsulin was excellent (Fig. 8B *left*), approaching 100% (Supplement Fig. S5). However, in the homozygote, little labeled insulin was produced (Fig. 8B), with a yield amounting to only 24% (Supplement Fig. S5). In a third experiment, recovery of mature insulin from newly-synthesized proinsulin in a euglycemic heterozygote again approached 100% (Fig. 8C), whereas in a homozygote with a random blood glucose of 537 mg/dL, insulin yield was only 4% (Supplement Fig. S5). Remarkably, in each case, newly-synthesized proinsulin in the homozygous *Ins2*-proinsulin-R(B22)E islets was detected by nonreducing SDS-PAGE as a ladder of aberrant disulfide-linked proinsulin complexes that exceeded the recovery of monomeric proinsulin. Indeed, recovery of newly-synthesized monomeric proinsulin by nonreducing SDS-PAGE worsened with the progression of hyperglycemia, tightly correlating increased proinsulin disulfide-linked complex formation with decreased insulin production.

**Fig. 8.**
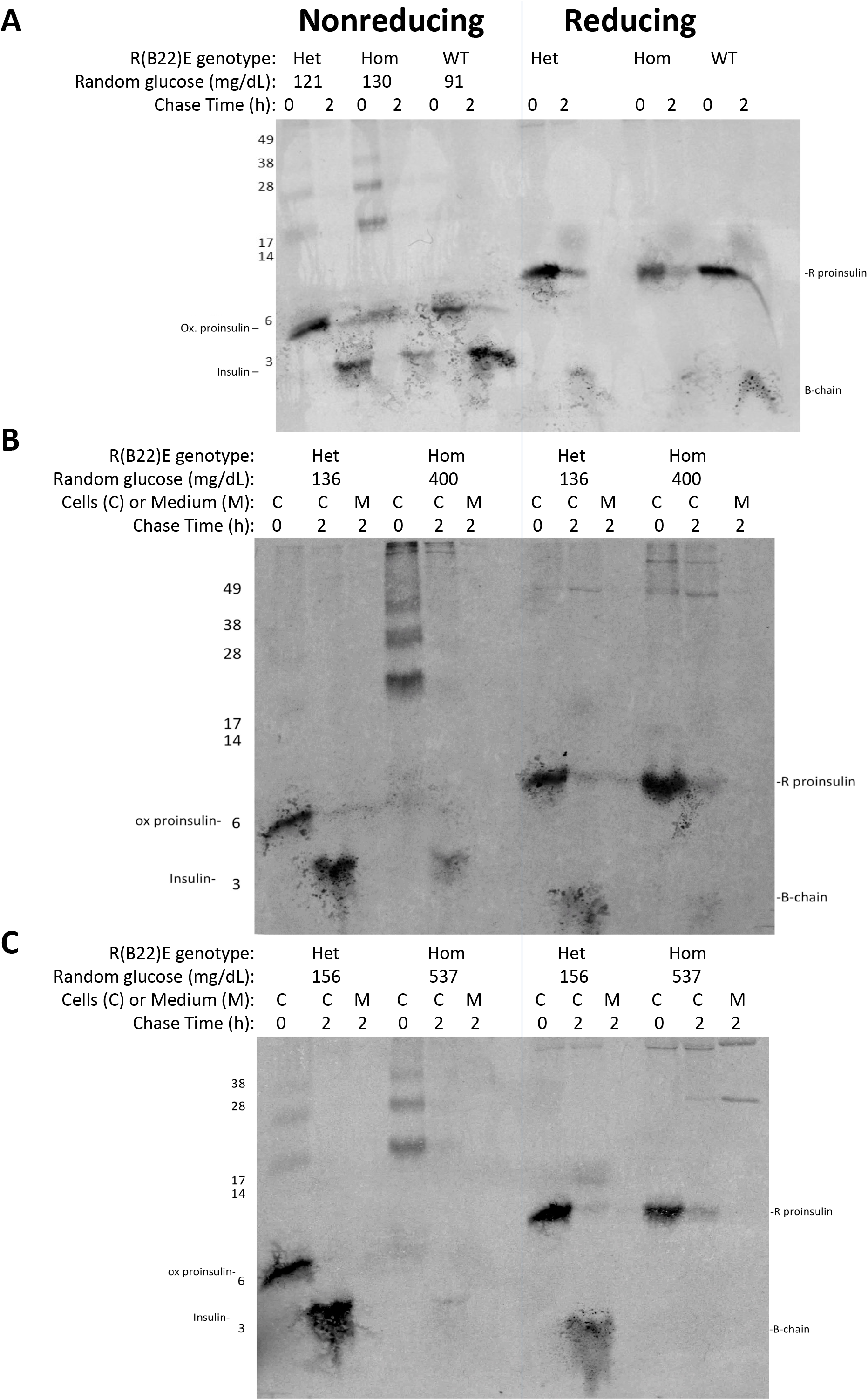
Biosynthesis of proinsulin and insulin in wild-type control and *Ins2*-proinsulin-R(B22)E heterozygous and homozygous males (with progression of diabetes, age 4-8 weeks). The random blood glucose of each animal at the time of euthanasia is indicated. Isolated islets were pulse-labeled with ^35^S-amino acids for 30 min and then chased for 2 h, as indicated. Cells (‘C’) and chase media (‘M’) were either combined together (so that no protein was lost) or analyzed separately. Samples were immunoprecipitated with anti-insulin followed by Tris-tricine-urea-SDS-PAGE under nonreducing (‘NR’, *left half of each gel*) or reducing conditions (‘R’, *right half of each gel*), followed by fluorography. A line is drawn separating the nonreduced and reduced samples, but these images show the complete gels and no lanes have been excised. **A, B and C** represent three independent experiments with the genotypes shown (ages 4, 5, and 8 weeks, respectively); the data are quantified in Supplement Fig. S5.

## Discussion

We report that unlike *Akita*-proinsulin-C(A7)Y, proinsulin-R(B22)E can pass ER quality control to become secreted from pancreatic β-cells (Fig. 1B), and its expression does not efficiently block export of co-expressed WT proinsulin (Fig. 1C). Moreover, in a normal laboratory environment, heterozygous *Ins2*-proinsulin-R(B22)E males gain weight post-weaning, exhibit normal glucose tolerance, and diabetes does not develop (Fig. 2A-D) for as long as we have followed the animals (up to 6 months of life). Nevertheless, proinsulin-R(B22)E does show evidence of misfolding in the ER (Figs. 1, 7, 8, Supplement Fig. S1). Additionally, when transitioned to a HFD, all male heterozygotes develop progressive hyperglycemia/diabetes (Fig. 2D-F) with inadequate insulin secretory response (Fig. 2G, H) due to diminished islet insulin content (Fig. 2J) accounting for impaired glucose-stimulated insulin secretion (Fig. 2I) despite the persistence of proinsulin (Fig. 2J) in a growing percentage of β-cells (Fig. 4B). A similar phenotype develops in homozygotes even without HFD (Fig. 3A-F; Fig 4C-E). An increase of circulating proinsulin-to-insulin ratio has been described in human T2D (34; 35); an intra-islet increase of proinsulin-to-insulin has also been reported during development of spontaneous T2D in animal models (36); and we also observe an increasing proinsulin-to-insulin ratio in the circulation (Fig. 3H) and in the islets themselves (Fig. 4C-E; Fig. 7A) in diabetic homozygotes.

Proinsulin misfolding is causal for diabetes in the homozygotes, with obviously aberrant disulfide-linked proinsulin complexes forming immediately upon synthesis (Fig. 7B, Fig. 8) and accompanying diminished insulin biosynthesis (Fig. 8, Supplement Fig. S5). Whereas the ability of isolated islets from WT *vs* heterozygous *vs* homozygous mice to respond to a glucose challenge exhibits no obvious defect in *fold-stimulation*, a progressive deficiency of insulin secretion under both unstimulated and stimulated conditions is apparent *in vivo* and *in vitro* (Fig. 2I, Figs. 3E, F) consistent with a failure to maintain the insulin storage pool (37).

The progression of dysglycemia/diabetes is linked to the emergence of intra-islet heterogeneity, with increasing proinsulin-rich cells (38-41) bearing little or no stored insulin — distinct from the subpopulation of β-cells rich in stored insulin (Fig. 4). Heterogeneity within the β-cell population is increasingly recognized at the mRNA level (42) including the description of “extreme” β-cells with lower insulin levels and higher juxtanuclear proinsulin immunostaining (43) — thought to represent the Golgi region from which new secretory granules emerge (23; 44). However, the diabetogenic progression in HFD-challenged heterozygous *Ins2*-proinsulin-R(B22)E mice (or homozygotes) involves islet cells exhibiting a proinsulin localization that fills the cytoplasm (Fig. 4) with expansion of the ER (Fig. 5, Supplement Figs. S3) and with increased generation of misfolded proinsulin (Figs. 7, 8) that co-localizes with KDEL-containing ER resident proteins (Supplement Fig. S4) accompanied by deficient insulin biosynthesis (Supplement Fig. S5). Curiously, this is what has been observed for the localization and misfolding of proinsulin in islets of T2D-like mice with hyperphagia-induced dysglycemia without any *Ins* gene variant (17). The total lifespan of proinsulin in β-cells is limited to ∼4 hours (23), suggesting that β-cells lacking insulin but bearing proinsulin [which may be entirely overlooked in β-cell mass measurements (45)] remain biosynthetically active right up to the time of our analysis.

Additionally, even in “off-scale high glucose” *Ins2*-proinsulin-R(B22)E homozygotes, there persists a subset of β-cells with substantial stored insulin content and only modest proinsulin (although they represent a shrinking fraction of total, Figs. 4, 6). Conceivably, such cells could represent ‘immature’ β-cells with deficient glucose-sensing (46; 47) that may not release stored insulin — but this remains to be determined. Additionally, the diseased islets develop increased glucagon-positive cells within the islet interior (Fig. 6), a feature noted in several diabetes models. Altogether, these data are consistent with the existence of islet β-cell subpopulations exhibiting heterogeneity in ER homeostasis (48) and an increase in islet “α-to β-cell” ratio (49).

In summary, the main observation in this manuscript is that proinsulin misfolding can be entirely subclinical, yet dramatic pathology emerges upon HFD exposure triggering rapid insulin deficiency. HFD-induced β-cell failure has been proposed to be alternatively associated with gluco/lipo-toxicity, β-cell senescence, de-differentiation, trans-differentiation, or apoptosis (50). This manuscript does not resolve those alternatives but highlights intraislet heterogeneity, with decreasing insulin-hi/proinsulin-low cells, increasing proinsulin-hi insulin-low cells, and glucagon-positive cells during disease progression. Unequivocally, our data in these models show that development of hyperglycemia runs anti-parallel with pancreatic insulin storage, i.e., biosynthesis of new insulin secretory granules is inadequate to replace the depletion of stored insulin that utilized to meet the body’s metabolic needs. Fascinating work is ongoing worldwide to understand the changes in islet cell heterogeneity during development of diabetes (48-50). We merely emphasize here that all of the pathological changes identified herein can be triggered by a genetic predisposition to proinsulin misfolding. Therefore we conclude that predisposition to proinsulin misfolding serves as an important potential risk factor to diet-induced diabetes.

## Acknowledgements

This work was supported by NIH R01-DK48280 and NIH P01 AI-118688. We acknowledge support from Leroux Devon in the Michigan Biomedical Research Electron Microscopy Core Facility, the Michigan Tissue and Molecular Pathology (Histology) Core for help with sample preparation, the Michigan Diabetes Research Center Morphology Core (NIH P30 DK020572), the University of Michigan Protein Folding Diseases Initiative, as well as institutional funds from the Howard Hughes Medical Institute at University of Colorado Anschutz Medical School, and National Jewish Health. Author contributions: PA and JK initiated and designed the research project; NJ collaborated with Dr. Jennifer Matsuda and the team in the National Jewish Mouse Genetics Core Facility in the design and creation of the original NOD knock-in mouse that gave rise to the mice used in these studies; DL developed analysis techniques; MA, AA, LH, NJ generated research data, reviewed data, and contributed to discussion; PA wrote the manuscript; all authors edited and reviewed the manuscript. Dr. Peter Arvan is the guarantor of this work and, as such, had full access to all the data in the study and takes responsibility for the integrity of the data and the accuracy of the data analysis.

**The authors have no conflicts of interest to declare**.

---

**Table I.**
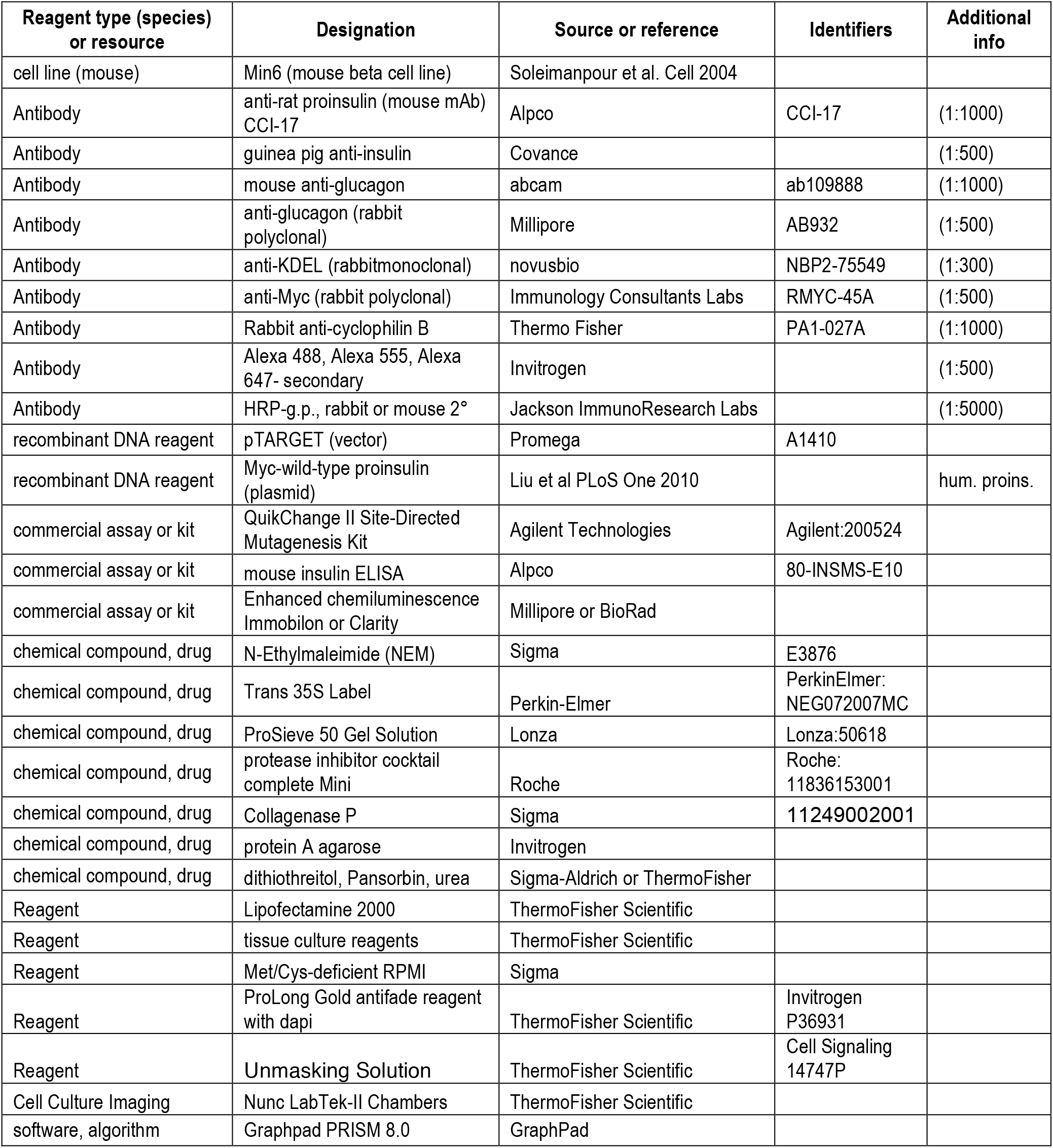
Key Resources.

