## Supplement file for "Predisposition to Proinsulin Misfolding as a Genetic Risk to Diet-Induced Diabetes"

**Supplement Figure legends**

**Supplement Fig. S1**. A portion of Figure 2J is reproduced (above); nonreducing SDS-PAGE and anti-proinsulin immunoblotting is shown below, highlighting the presence of aberrant disulfide-linked proinsulin complexes.

**Supplement Fig. S2**. Circulating proinsulin, and proinsulin-to-insulin ratio, from 11.5 week old HFD-fed *Ins2*-proinsulin-R(B22)E heterozygous males (pink symbols), superimposed onto the data reproduced from Figs 2G and H.

**Supplement Fig. S3**. Transmission electron micrographs of islets cells from the genotypes (and diet) included in this study. Random blood glucose at the time of euthanasia is indicated on each figure. Scale bars are shown on all figures. **A)** Typical β-cell from male WT mouse and *Ins2*-proinsulin-R(B22)E heterozygote on normal chow diet (age 4 weeks). **B)** Another typical β-cell from the animals analyzed in panel A, at higher magnification to highlight ER and secretory granule morphology. **C)** β-cell from WT mouse fed a HFD for 6 weeks with organelles labeled on the figure. ISG = immature secretory granule. VTCs are a pre-Golgi compartment. **D)** *Upper left image*: low-power of islet from HFD-fed male *Ins2*-proinsulin-R(B22)E heterozygote (age 6 weeks). "Cell A and B" are β-cells; "Cell C" is an α-cell. Note that Cell B is poorly granulated. *Right*: A magnified image from the white-boxed region of the upper left micrograph, showing that Cell B has expanded ER and numerous small and under-filled insulin secretory granules with low electron-density contents. *Lower left*: A magnified image from the green-boxed region of the upper left micrograph, showing a neighboring portion of the cytoplasm of Cell B with numerous small and under-filled insulin secretory granules. **E)** *Ins2*-proinsulin-R(B22)E male homozygote (age 4 weeks) highlighting a β-cell essentially lacking all secretory granules but with markedly expanded ER. A fragment of the cytoplasm of a neighboring cell bearing secretory granules is visible in the lower left corner. **F)** A portion of the cytoplasm of two further β-cells from the animal in panel E: the upper cell shows a profusion of ER with very few micro-granules; the lower cell shows a profusion of under-filled micro-granules (i.e., abnormally small size and lower-than-normal electron density of contents). **G)** Several further β-cells from the animal in panel E; a portion of the cytoplasm of an α-cell is shown at the bottom right. The main β-cell captured almost in its entirety, albeit uncommon, highlights expanded ER and under-filled secretory pathway organelles; the cytoplasm of other β-cells in the lower left corner or upper right corner of the same image do show sparse insulin granules. *Boxed image at right*: A magnified image from the red-boxed region of the left micrograph, highlighting under-filled secretory pathway organelles with low electron-density contents.

**Supplement Fig. S4**. ER resident proteins (anti-KDEL, blue) in subpopulations of proinsulin-enriched cells (red) and insulin-enriched cells (green), as identified by triple immunofluorescence (merged images shown). The genotypes and random blood glucose at the time of euthanasia is noted on each image. Note that KDEL proteins are widely expressed in the exocrine pancreas (blue), which lacks proinsulin or insulin. A purple merged image derives from the sum of proinsulin (red) and KDEL proteins (blue). The animal genotypes are shown on the figure (females, age 5-6 weeks).

**Supplement Fig. S5.** Recovery of newly-synthesized insulin derived from pulse-labeled proinsulin, derived from the phosphorimages in Fig. 8; genotype and random blood glucose as indicated on the figure. Proinsulin bands (at chase time zero) were quantitated from reducing gels;

newly-synthesized insulin derived from the pulse-labeled samples were quantified from nonreducing gels (insulin is a two-chain protein which “falls apart” under reducing conditions; thus nonreducing gels are preferable for this analysis).

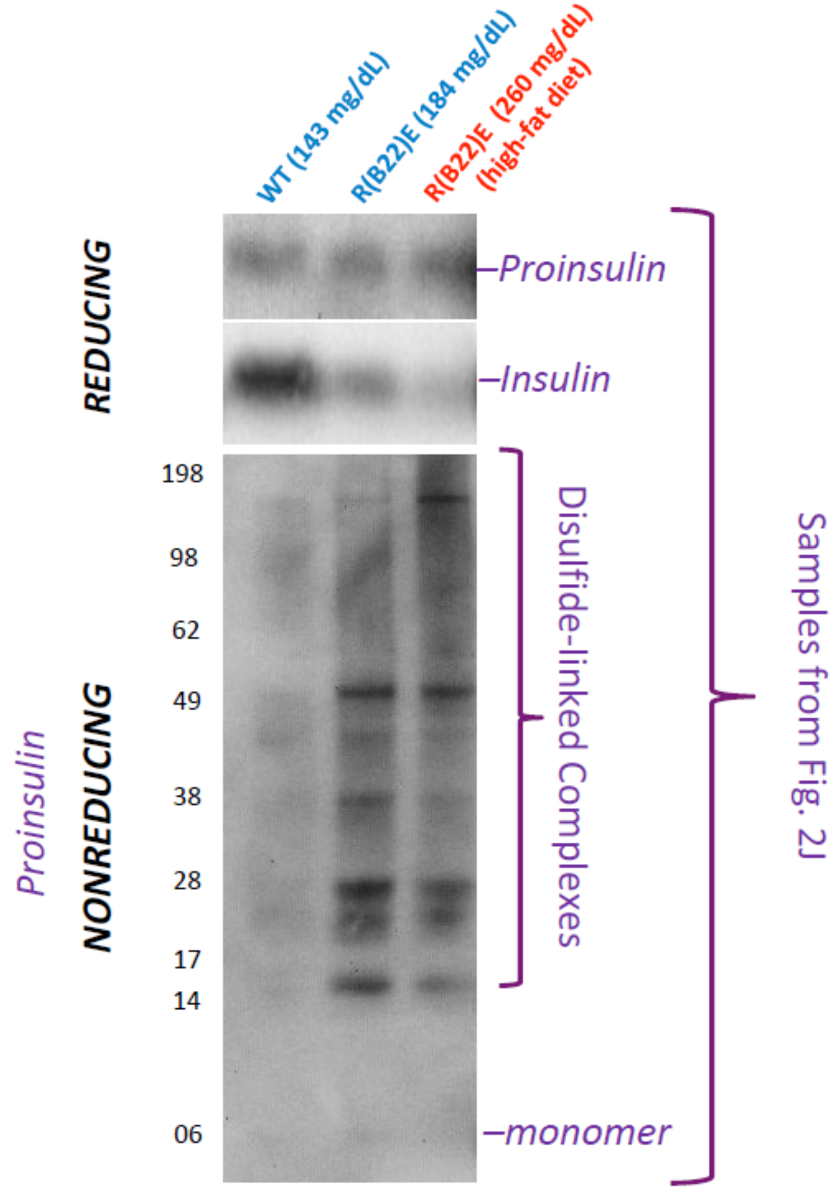


Maroof et al., Fig. S1

R(B22)E-Het (Male)HFD mice included in these graphs

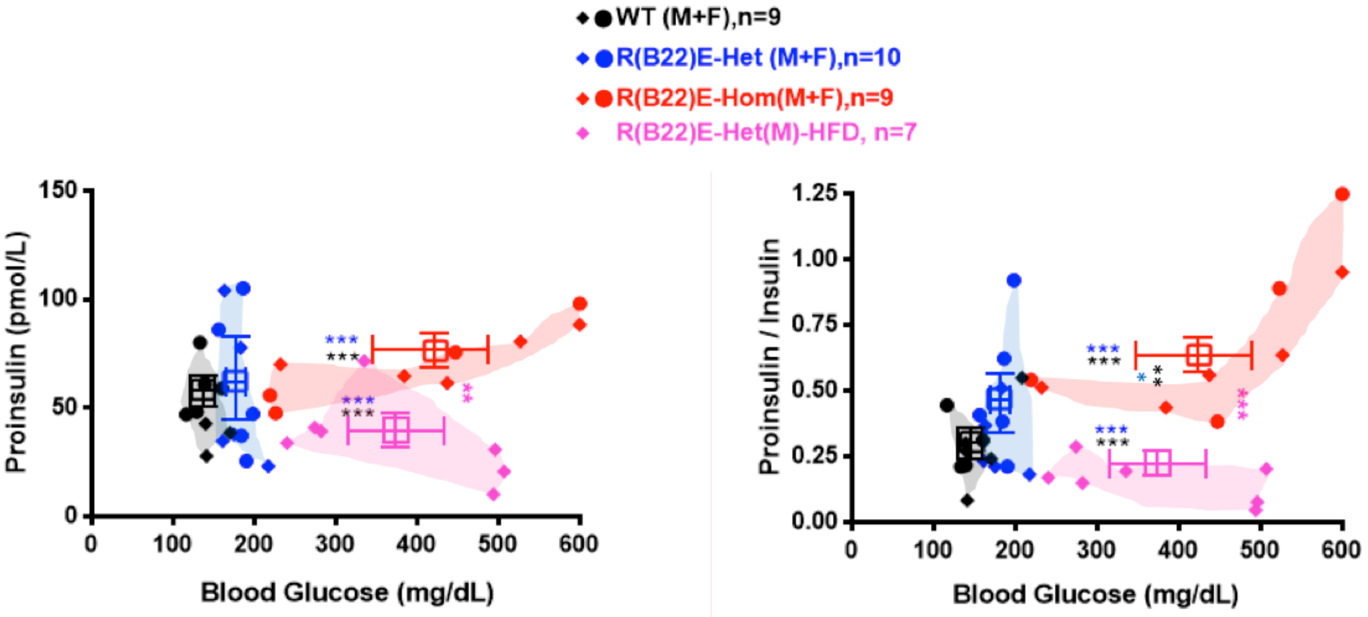


**For proinsulin**
WT vs Hom, P=0.14
Het(NC) vs Hom, p=0.26
WT vs Het(NC), p=0.98
WT vs Het(HFD), p=0.34
Het(NC) vs Het(HFD), p=0.19

**For proinsulin / Ins**
WT vs Het(NC), p=0.63
WT vs Het(HFD), p=0.59
Het(NC) vs Het(HFD), p=0.09

Maroof et al., Fig. S2

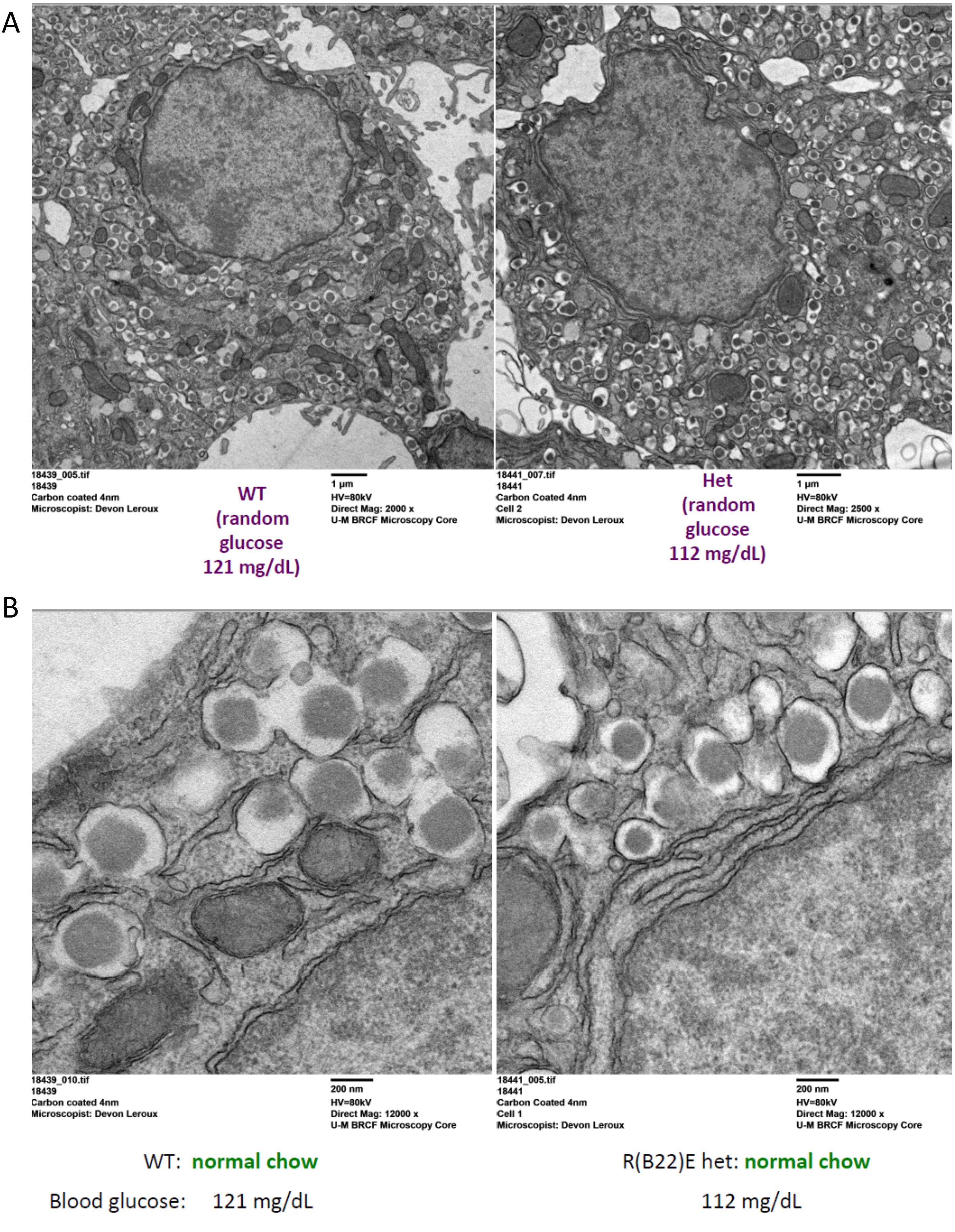


Maroof et al., Fig. S3

C

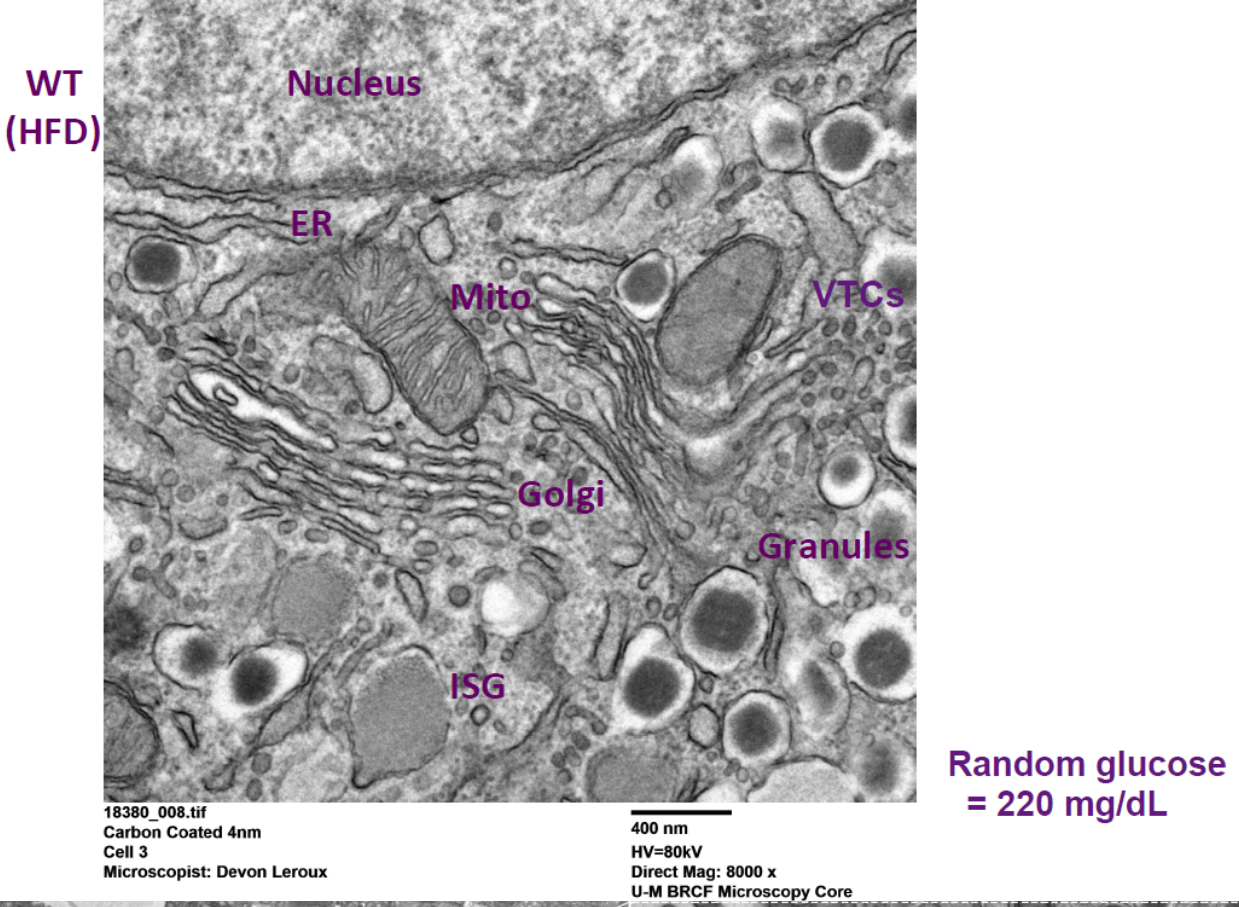


D

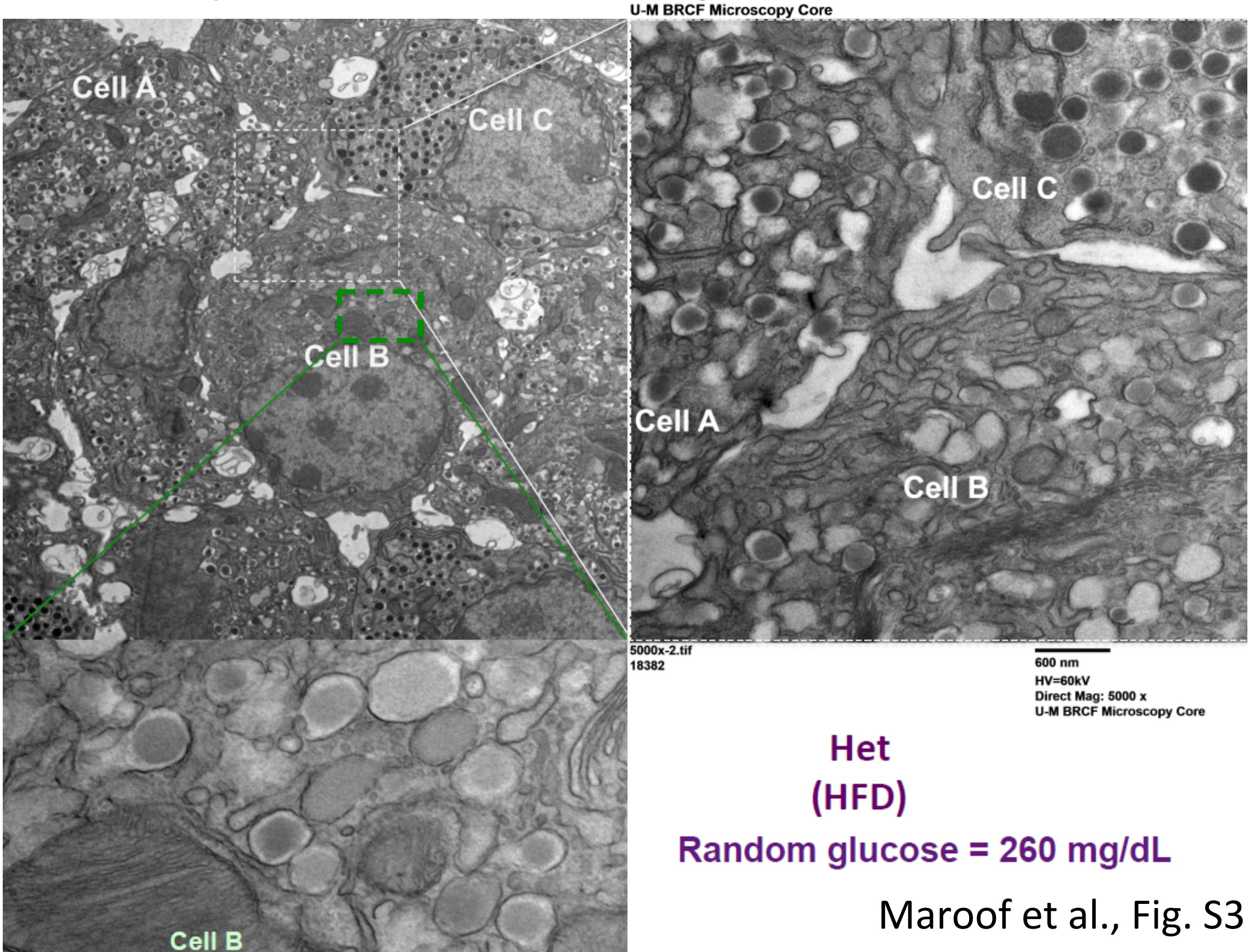


Maroof et al., Fig. S3

E

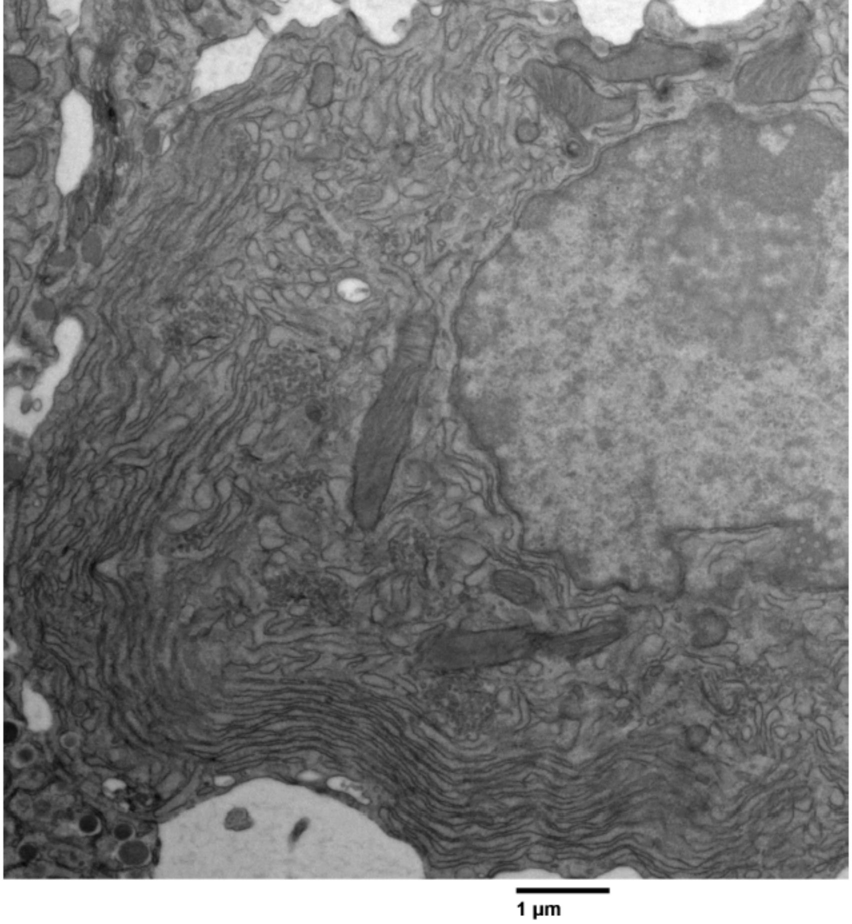


F

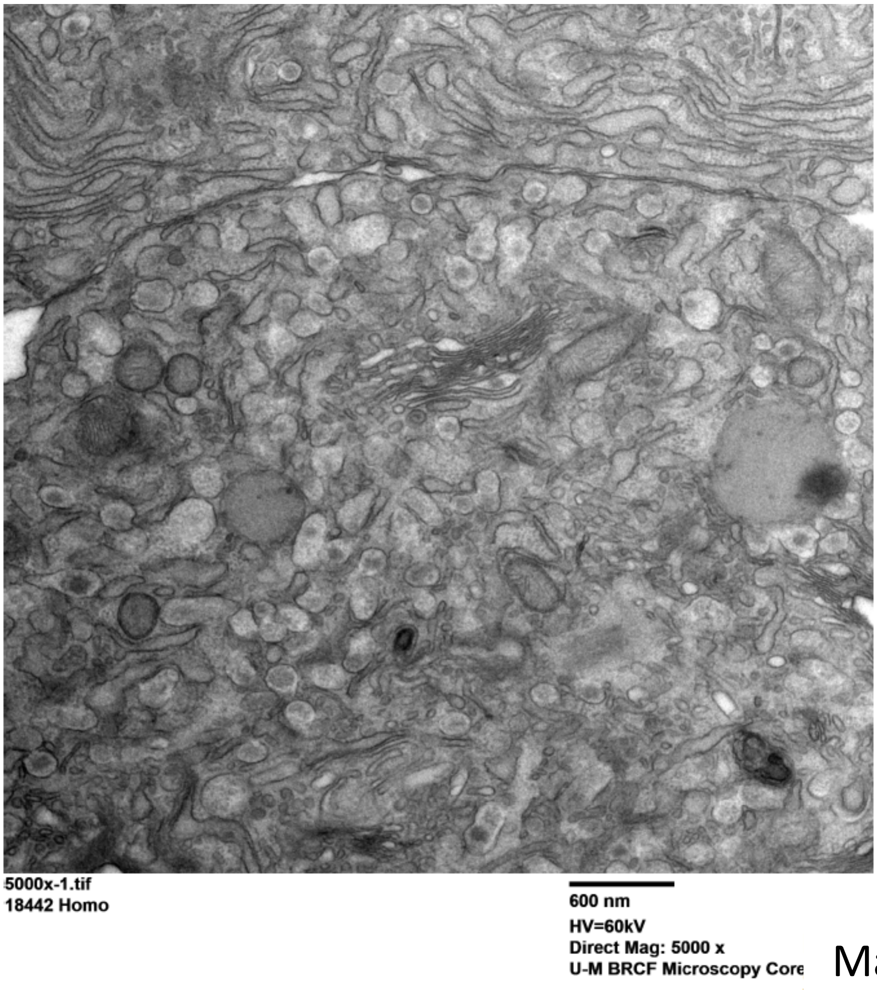


Maroof et al., Fig. S3

G

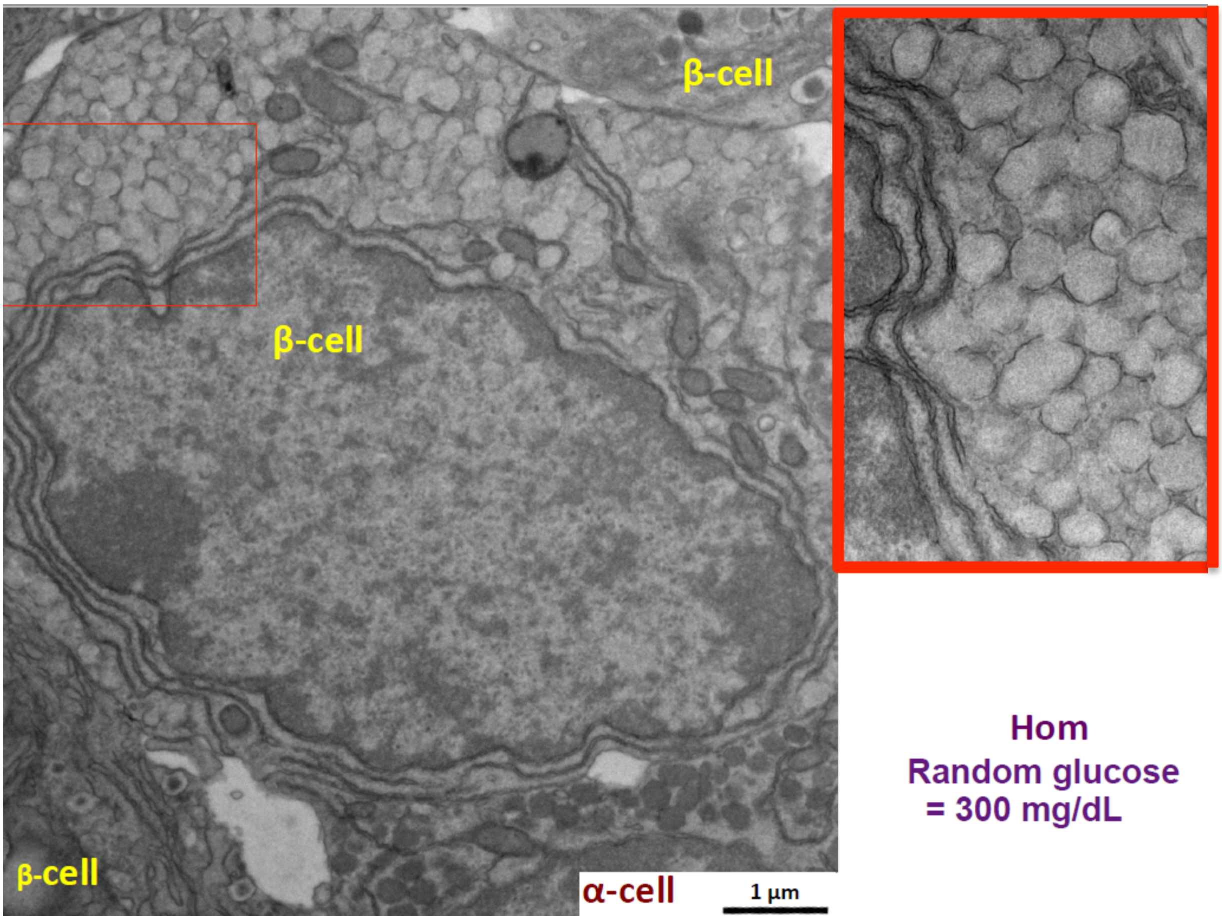


Hom
Random glucose
= 300 mg/dL

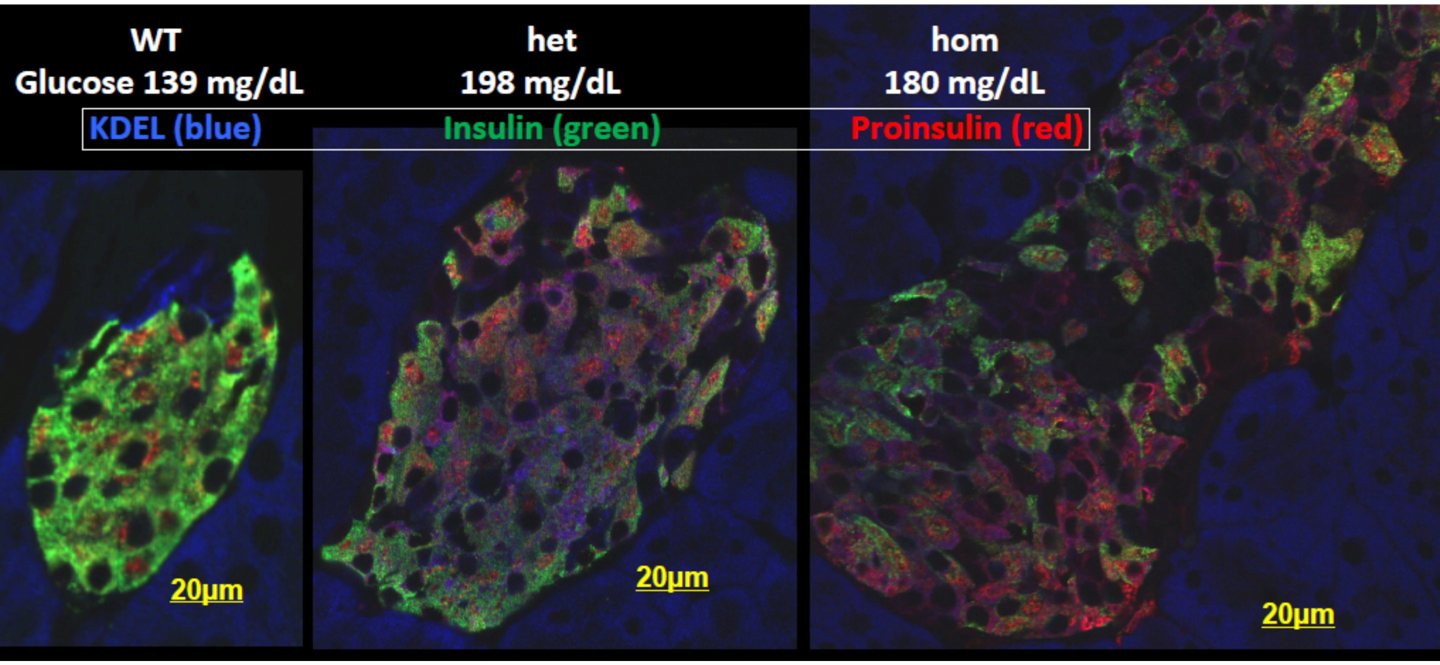


Maroof et al., Fig. S4

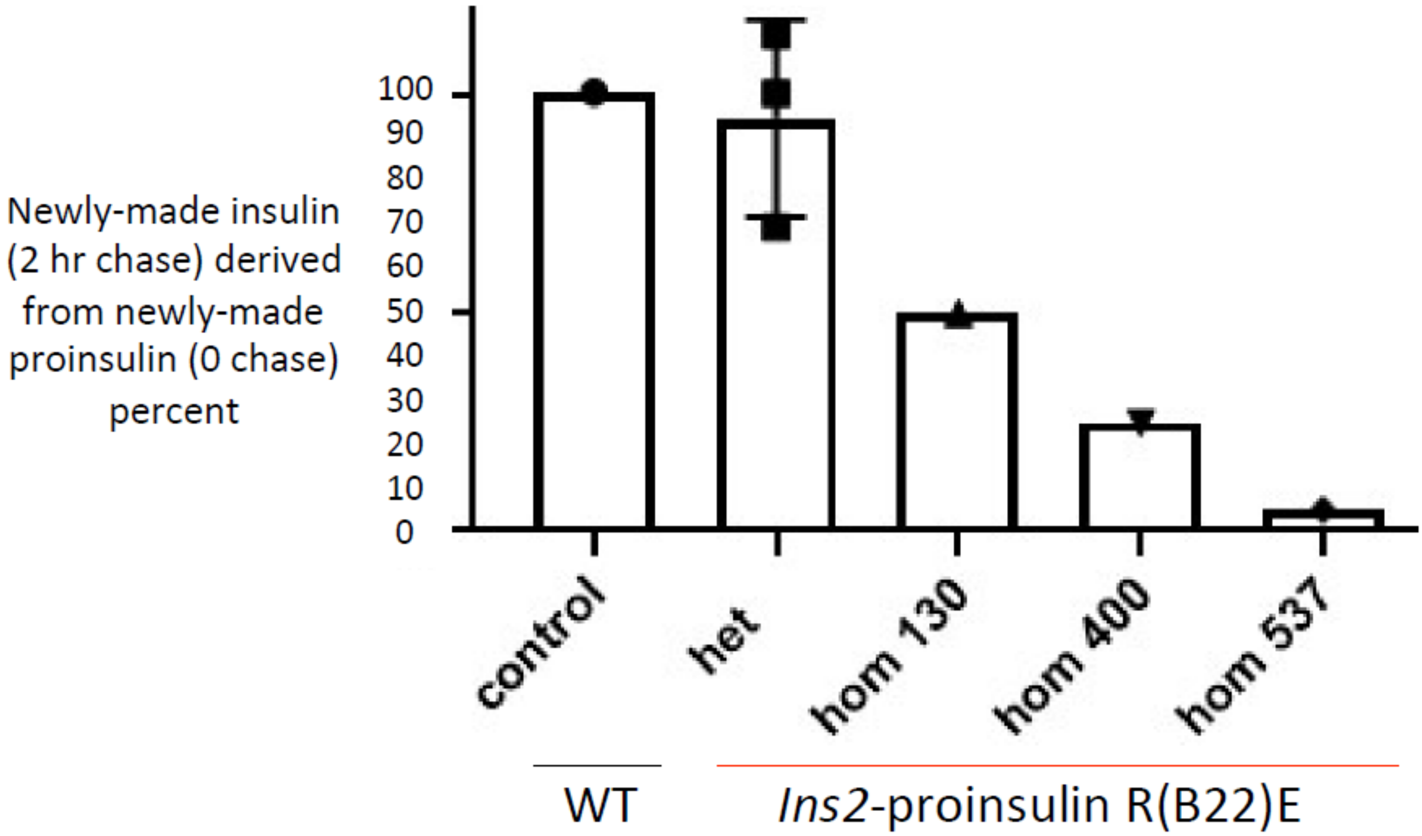


Maroof et al., Fig. S5
